# Missense mutations in intrinsically disordered protein regions link pathogenicity and phase separation

**DOI:** 10.1101/2025.02.19.639140

**Authors:** Oliver L. Kipp, Karen A. Lewis, Loren E. Hough, Steven T. Whitten

## Abstract

The impact of missense genetic variations on protein function is often enigmatic, especially for mutations that map to intrinsically disordered regions (IDRs). Given the functional importance of phase separation of IDRs, it has been proposed that mutations that modulate phase separation might preferentially lead to disease. To examine this idea, we used the robust predictability of phase-separating (PS) IDRs and annotation of disease-associated proteins and mutations to map the correlation between disease and phase separation. Consistent with previous work linking phase separation to cancer and autism spectrum disorder, we find a higher prevalence of predicted phase separation behavior in disease-associated proteins than typical for human proteins. We map the prevalence of phase separation across a wide range of diseases, finding that many, but not all, show an enrichment of phase separation in the proteins associated with them. Strikingly, the pathogenic mutation rate in predicted PS IDRs was elevated three-fold relative to IDRs not predicted to phase separate. Substitutions involving arginine and the aromatic types were among the most pathogenic for PS IDRs, while substitutions involving serine, threonine, and alanine the most benign. We applied these trends to mutations of uncertain clinical significance and predict that half found in PS IDRs are likely pathogenic. We find that phosphorylation sites were enriched in PS IDRs when compared to other protein regions, though mutations at such sites were mostly benign. Pathogenicity was highest for mutations in predicted PS IDRs when also found in a short linear motif, known mediators of protein-protein interactions.

## Introduction

Large-scale sequencing efforts have resulted in catalogs of the human genome and its variations, providing insight on the genetic basis of traits and diseases (1–4). Genetic variations that cause single amino acid missense mutations are often found in the intrinsically disordered regions (IDRs) of proteins, as IDRs comprise approximately one-third of the human proteome (5, 6). IDR sequences are less conserved relative to the sequences found in protein regions that fold (7, 8). Disease-linked mutations, however, are more prevalent in the folded regions (9). The functional impact of mutations in folded regions often can be understood by their perturbations to structure and stability (10–12). For IDRs, disease-linked mutations have been found to disrupt protein-protein interactions (13, 14), change the sequence’s propensity for disorder (9), alter the dimensions (e.g., compaction) of the disordered conformational ensemble (15), and modify sites of post-translational modifications (14) or short linear motifs (SLiMs) (16).

IDRs have been associated with many key biological processes, including signal transduction (17, 18), chromatin remodeling (19), cell cycle control (20), and directing the curvature of membrane surfaces (21). IDRs also are essential mediators of liquid-liquid phase separation (22, 23), a phenomenon in which condensed liquid droplets separate from and subsequently coexist within bulk cytosol (24). Biological condensates such as nucleoli, Cajal bodies and stress granules are critical for intracellular compartmentalization and functional specificity (24–26). Human genetic variations that mutate IDR sequences thus potentially alter the function and properties of biological condensates, and possibly contribute to disease phenotypes (27).

Several studies provide evidence that dysregulation of protein phase separation has a role in human disease (28–30). Mittag and colleagues demonstrated that cancer-associated mutations in the tumor suppressor SPOP disrupt phase separation *in vitro* and interfere with the co-localization of SPOP with its substrates in nuclear speckles (31). Fusion oncoproteins, e.g., EWS-FLI1 (32, 33), spontaneously form condensates inside cells that promote cancer development (34, 35). Many proteins known to phase separate also have been found in neurodegenerative inclusions (30), for example ⍺-synuclein in Parkinson’s disease (36), tau in Alzheimer’s disease (37), and TDP-43 (38) or FUS (39) in amyotrophic lateral sclerosis. Additionally, several predictors of phase separation have been used to identify correlations between phase separation and disease (40, 41), including cancer and autism spectrum disorder (27).

Previously, we developed an algorithm, ParSe, that accurately identifies from the protein primary sequence those IDRs likely to exhibit physiological phase separation behavior (42, 43). ParSe uses an optimal set of amino acid property scales for fast predictions of domain-level structure and provides a simple, quantitative metric for the sequence-calculated phase separation potential that obtains reasonable predictive power for existing mutant data (43). ParSe shows similar accuracy to other methods that have been developed to predict which protein sequences drive phase separation (43–49). One advantage of the ParSe algorithm is its speed, which enables it to identify drivers of phase separation at proteome scale (50).

Here, we investigate if ParSe can be used to identify relationships linking protein-mediated phase separation, human disease, and genetic missense variations. We found that human proteins associated with disease were enriched in predicted phase separation behavior relative to human proteins in general. Moreover, the incident rate of confirmed pathogenic missense mutations was three times higher in predicted phase-separating (PS) IDRs compared to IDRs not predicted to phase separate. This increase was not observed for confirmed benign missense mutations, suggesting the pathogenicity of mutations found in predicted PS IDRs was linked to the predicted phase separation behavior rather than some other IDR-driven function. Missense mutations at sites found in a SLiM were especially pathogenic when also in a predicted PS IDR. In contrast, missense mutations at phosphorylation sites in a predicted PS IDR were primarily benign. Comparing pathogenic and benign mutations in predicted PS IDRs revealed that single amino acid substitutions involving arginine and tyrosine were among the most pathogenic, which agrees with multiple reports showing the importance of arginine and tyrosine content in driving protein-mediated phase separation (46, 51–54). However, the types of substitutions that tended to be pathogenic in predicted PS IDRs also tended to be pathogenic in the other protein regions as well. Collectively, our findings yield insight into the impact of genetic missense variations that mutate IDRs, revealing that pathogenicity associates with sequence features predicting IDR-mediated phase separation behavior.

## Results

### Construction of datasets

We first identified disease and comparison datasets to investigate the sequence-based differences between proteins in general and the subset that have been associated with disease, and between pathogenic missense mutations and benign missense mutations. The reference proteome for *Homo sapiens* (20,435 sequences) from UniProt (55) was used for the comparison set. Disease-associated proteins were obtained by searching UniProt for proteins annotated as “Human”, “Reviewed (Swiss-Prot)”, and the keyword “Disease”. Disease-associated proteins (4,639 sequences) were sub-grouped according to condition, e.g., age-related macular degeneration, amyloidosis, asthma. Human proteins (1,248 sequences) with entries in DisProt (56) were used to represent IDPs in general. Sequence sets representing phase separation exhibited varying levels of curation: human proteins (3,917 sequences) annotated as “condensate” in CD-CODE (57), human proteins (129 sequences) annotated as “condensate driver” in CD-CODE, and *in vitro* confirmed phase-separating human proteins (66 sequences) assembled previously (42) from Vernon et al (46), DisProt (56), and PhaSePro (58).

Confirmed pathogenic and benign missense mutations were obtained from two sources. First, we used the humsavar index of human variants curated from literature reports, which is part of the UniProt Knowledgebase (55). This index contained 32,665, 39,656, and 10,178 single amino acid substitutions that have been classified as likely pathogenic or pathogenic, likely benign or benign, or of uncertain clinical significance, respectively. We constructed a second missense mutation set from a natural variants index from UniProt. Mutations in this index were imported from the Ensembl Variation (59) and ClinVar databases (4) and include human variations from the 1,000 Genomes Project (3, 60). A small percentage (3%) of the single amino acid missense mutations in this index included predicted clinical significance; 59,435, 42,617, and 919,968 were predicted as likely pathogenic or pathogenic, likely benign or benign, or of uncertain clinical significance, respectively, and 35,019,669 single amino acid substitutions had no predicted clinical significance.

Confirmed SLiMs, which are short stretches of protein sequence that mediate protein-protein interactions, were obtained from the Eukaryotic Linear Motif (ELM) resource (61). Search of the ELM database for all “instances” filtered for “true positive” and “Homo sapiens” found 2,247 SLiMs from 1,371 human proteins. A curated index of 531,023 experimentally confirmed protein phosphorylation sites from 30,637 human proteins was obtained from the Eukaryotic Phosphorylation Site Database 2.0 (62).

### Sequence-based calculation of phase separation potential

Dysregulation of protein phase separation has been associated with several human diseases (27–30). To explore this relationship broadly, we used the ParSe algorithm (43) to examine how different types of mutations (pathogenic, benign, or of unknown clinical significance) correlate with phase separation. Additionally, we compared the predicted prevalence of phase separation of disease-associated proteins to the reference human proteome. ParSe predicts the domain-level organization of a protein from the primary sequence; for folded, ID, and PS ID. The prediction is based on our finding of robust property differences between the three classes of protein regions (43). As diagramed in Figure S1, we use a sliding-window approach to find where local properties in a sequence match the folded, ID, and PS ID classes. Figure 1 shows the ParSe prediction for multiple human proteins confirmed to phase separate *in vitro* (51, 63–68).

**Figure 1.**
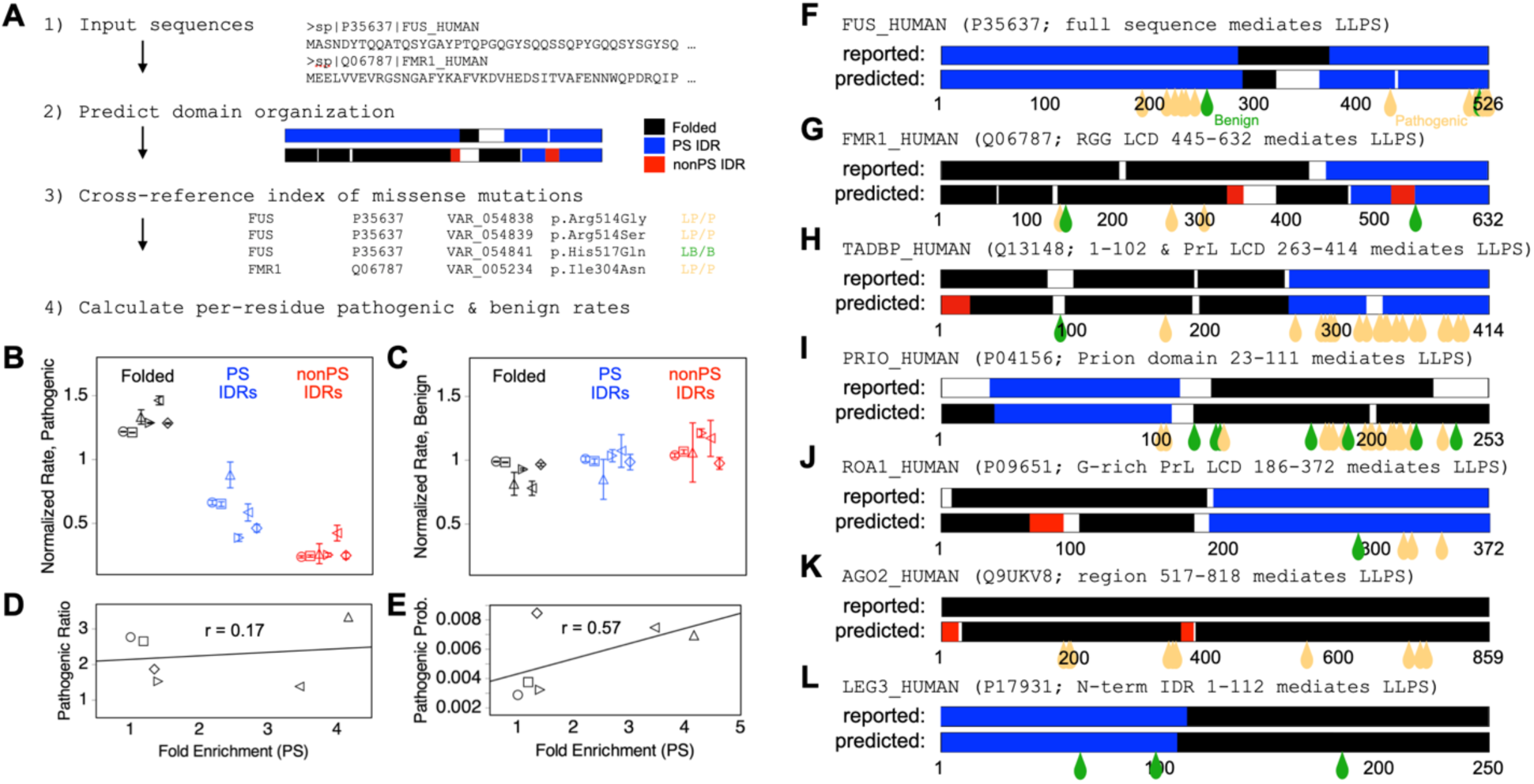
Pathogenic and benign mutation rates by protein region. (**A**) Steps used to calculate mutation rates. Domain organization was predicted from sequence by ParSe. Pathogenic and benign missense mutations were obtained from the humsavar index. Per residue (**B**) pathogenic and (**C**) benign mutation rates by region class were calculated for, from left to right, the reference human proteome (○), proteins with entries in humsavar (□), *in vitro* verified phase-separating human proteins (△), human proteins annotated condensate in CD-CODE (▷), human proteins annotated condensate driver in CD-CODE (◁), and human proteins with entries in DisProt (◇). Error bars show the standard error. (**D**) Sequence sets curated for phase separation had the highest enrichment for proteins predicted PS relative to the reference proteome (“Fold Enrichment (PS)”); the ratio of pathogenic mutation rates (“Pathogenic Ratio”), PS IDR rate divided by nonPS IDR rate, correlated weakly with the fold enrichment (r = 0.17). (**E**) Per residue probability of finding pathogenic mutations exhibited a correlation with the fold enrichment (r = 0.57). Reported versus predicted domain organization for (**F**) FUS, (**G**) FMRP, (**H**) TDP-43, (**I**) PrP, (**J**) hnRNP A1, (**K**) AGO2, and (**L**) Galectin-3. Teardrops show sites of pathogenic (light orange) and benign (green) missense mutations.

ParSe also can be used to calculate a predicted phase separation potential for individual proteins, taken as the sum of the classifier distance of each window matching the PS ID class (Figure S1). The classifier distance was developed to assess confidence in the window assignments (43). A higher “potential” can arise from longer predicted PS IDRs, a greater number of predicted PS IDRs, and higher confidence in these predictions. To enable a direct comparison between protein sets, we considered a cutoff in PS potential of 100. This value was chosen because every protein in a set of 43 confirmed to exhibit homotypic phase separation behavior, curated by Vernon et al (46), had a computed PS potential of at least 100 (Figure S2). We use that minimum potential as a cut-off throughout this paper, identifying PS predicted proteins as those with a PS potential above 100.

### Mutations in IDRs link pathogenicity and phase separation

We sought to compare rates of missense mutations in PS IDRs and nonPS IDRs (Figure 1). Strikingly, we found significant enrichment of pathogenic mutations within PS IDRs as compared to nonPS IDRs. Normalized mutation rates were calculated as the proportion of mutations divided by the proportion of residues in a region class (Tables S1 and S2). Figure 1B shows that the normalized rate for pathogenic missense mutations in predicted PS IDRs (0.66 ± 0.02, rate ± standard error) was approximately half the rate found in predicted folded regions (1.22 ± 0.002). The PS IDR rate was almost 3-fold higher than the rate calculated in predicted nonPS IDRs (0.24 ± 0.009). Thus, though the normalized rate for pathogenic missense mutations was highest in folded protein regions, pathogenic missense mutations in IDRs were primarily found in IDRs with sequence features predicting phase separation behavior. The pathogenic versus benign odds ratio also revealed that folded regions exhibited a 2-fold higher pathogenicity compared to PS IDRs, while PS IDRs had a 3-fold higher pathogenicity than nonPS IDRs (Figure S3A).

This ranking of pathogenic mutation rates by region class (folded > PS ID > nonPS ID) was observed in subset proteomes of proteins with confirmed mutations, confirmed IDRs, and confirmed phase separation behavior (Figure 1B). The sets curated for phase separation showed increases in fold enrichment for PS compared to the sets curated for ID and mutation (Figure 1D). Fold enrichment was calculated as the fractional number of PS predicted proteins in a set divided by its fractional number in the reference human proteome. The ratio of pathogenic mutation rates, PS IDR rate divided by nonPS IDR rate, was >1 for each sequence set and correlated weakly with fold enrichment for PS (r = 0.17), suggesting that bias from the computational predictor, or that could arise from some proteins being more researched than others, was minimal. Moreover, the probability of finding a pathogenic missense mutation in a sequence set (mutations found divided by total residues) increased for sets curated for confirmed phase separation behavior (Figure 1E). This shows that mutations in PS proteins are more pathogenic.

In contrast, there was no significant difference in the normalized rate by protein region class for benign single amino acid missense mutations, found to be ∼1 in predicted PS ID, nonPS ID, and folded protein regions, for each of the full reference and subset proteomes (Figure 1C). Figures 1F-L show the reported and predicted domain-level organization of representative proteins *in vitro* verified to phase separate (51, 63–68), as well as the sites of their confirmed pathogenic and benign missense mutations. Qualitatively similar results were obtained with the natural variants index, whereby the normalized rate for pathogenic missense mutations was highest in protein regions predicted to be folded, followed by PS IDRs, and then substantially lower in nonPS IDRs (Figure S4).

We then sought to determine if these domain-specific differences in pathogenic mutation rates could be generalized to proteins containing significant PS regions and found that all mutations are overrepresented in PS predicted proteins as compared to the human proteome. Using a cutoff of PS potential of 100, ∼16% of the human proteome and ∼21% of the proteins with pathogenic mutations, a 1.3-fold enrichment, are predicted to exhibit homotypic phase separation behavior (Figure S5). The distribution of PS potential values in the two sets showed a statistically significant difference by the nonparametric Mann-Whitney *U*-test (69), giving a one-tail *p*-value < 2.2e-16. Similar results were obtained for proteins with benign or of uncertain clinical significance mutations in the humsavar index, showing 1.2-fold and 1.6-fold enrichment, respectively. Qualitatively similar results were obtained with the natural variants index, whereby proteins harboring single amino acid missense mutations were enriched in predicted phase separation behavior (Figure S5). Thus, proteins with sequence features predicting phase separation were more likely to have identified mutations. This may be due to the differences in overall size and composition, as nonPS proteins (predicted PS potential below 100) were, on average, shorter in length and more folded as compared to PS predicted proteins (Figure S6).

### Mutations in SLiMs within PS IDRs exhibit exceptionally high pathogenicity

SLiMs are conserved recognition motifs involved in signal transduction (70) and found primarily in the IDRs of the human proteome (71). As protein-protein interactions are fundamental to the functional outcomes of phase separation, enabling the formation, regulation, and biological specificity of biomolecular condensates (26, 72), we next asked if proteins containing SLiMs were enriched for phase separation. Using ParSe, we found that 38.1% of human proteins with confirmed SLiMs were predicted to exhibit phase separation behavior; a 2.3-fold enrichment relative to the reference human proteome, where only 16.4% of human proteins are predicted to exhibit phase separation behavior (Figure S5).

Among IDRs, confirmed SLiMs were enriched more so in nonPS than PS IDRs. Here, the normalized rates were calculated as the proportion of confirmed SLiMs divided by the proportion of residues found in a region class. The computed rates in PS ID, nonPS ID, and folded regions were 1.8 ± 0.11, 2.5 ± 0.10, and 0.42 ± 0.01, respectively (Figure 2A).

**Figure 2.**
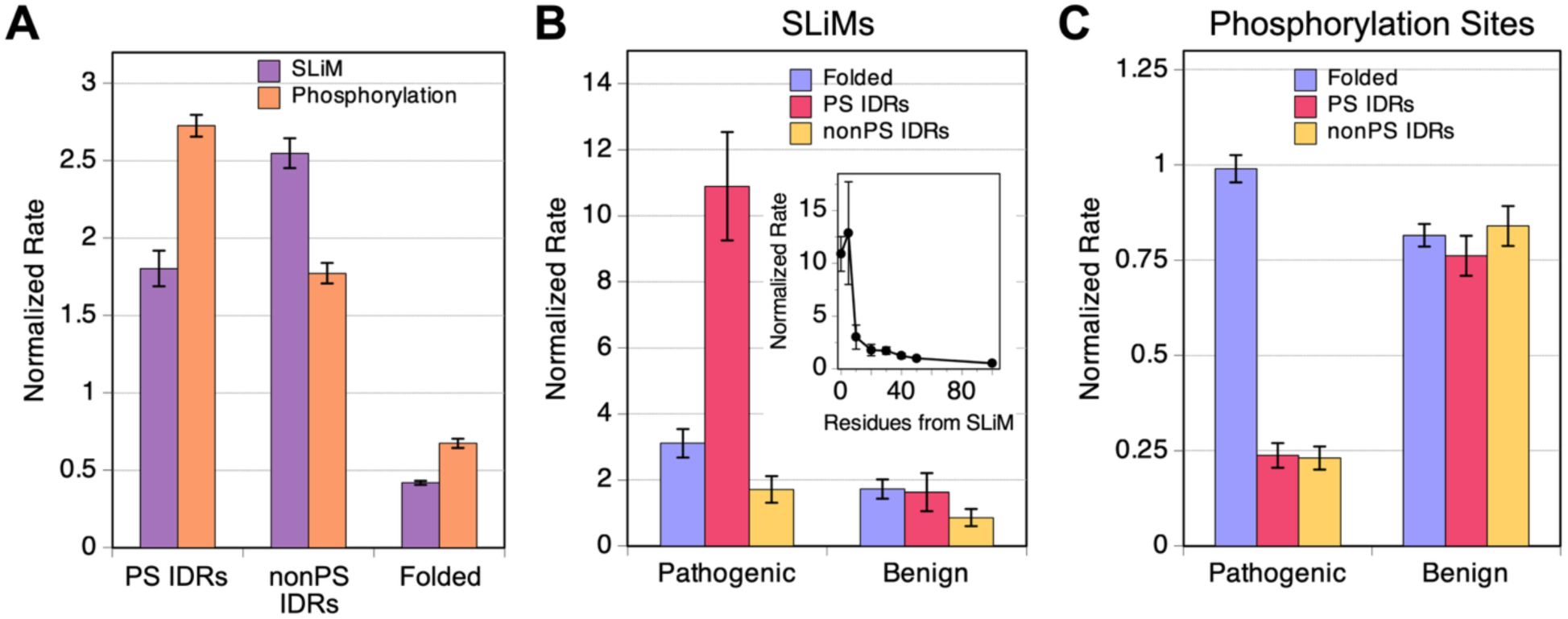
SLiM and phosphorylation rates by protein region and their rates of pathogenic and benign mutation. (**A**) Rates for finding a SLiM (purple) or phosphorylation site (salmon) by protein region class. Pathogenic and benign mutation rates for (**B**) SLiMs or (**C**) phosphorylation sites by protein region class. The panel **B** inset shows the pathogenic mutation rate for residues at positions located immediately outside the SLiM. Rates were calculated using the reference human proteome.

We next sought to quantify rates of missense mutations in SLiMs. Only 0.43% of confirmed pathogenic single amino acid missense mutations were found in a confirmed SLiM. However, because confirmed SLiMs were an even smaller fraction of the reference human proteome (0.14%), the computed normalized rate for pathogenic missense mutations in SLiMs was 3.03 ± 0.25. The normalized rate for confirmed benign missense mutations in a confirmed SLiM was 1.35 ± 0.15 (Table S2). This result matches the previous finding that, in IDRs, pathogenic missense mutations are more likely to occur within SLiMs than benign missense mutations (16).

Pathogenic mutations were particularly enriched in SLiMs in PS IDRs, though the overall number of such mutations was small. The normalized rate of pathogenic missense mutations in SLiMs in PS IDRs was 10.9 ± 1.6, the highest of all pathogenic rates found in this work; 44 of 32,665 pathogenic missense mutations (0.13%) in the humsavar index were found in both a confirmed SLiM and a predicted PS IDR, whereas confirmed SLiMs in predicted PS IDRs represented 0.012% of the reference human proteome (Figure 2B). For comparison, the normalized rate was 1.6 ± 0.6 for benign missense mutations found in both a confirmed SLiM and a predicted PS IDR. The pathogenic versus benign odds ratio for missense mutations in a SLiM in a predicted PS IDR was 5.6 ± 2.2 (Figure S3B), reflecting the significantly higher normalized pathogenic rate. The normalized rate for pathogenic mutation in both a SLiM and predicted nonPS IDR, and a SLiM and predicted folded region was 1.7 ± 0.4 and 3.1 ± 0.4, respectively (Figure 2B). Thus, pathogenicity increased for SLiM positions in each region class (Figure S7) relative to the pathogenic mutation rate that included nonSLiM positions (Figure 1A). However, pathogenicity was highest, and by a significant margin, for missense mutations in a SLiM that also occurred in a predicted PS IDR (Figure 2B). As such, even though SLiMs are more likely to be found in nonPS IDRs than PS IDRs, their mutation was more likely pathogenic in PS IDRs.

Notably, the computed pathogenic mutation rate drops to the base PS IDR rate (∼0.6 ± 0.1) with increasing residue distance from the SLiM (Figure 2B inset). Thus, the exceptionally high pathogenic missense mutation rate was concentrated on the SLiM in the IDR and not the nonSLiM positions. As conserved recognition sites, SLiMs and other protein-protein interaction interfaces facilitate biological phase separation (73) through various key roles, e.g., condensate formation, stability, internal structure, and composition, which may explain their exceptionally high mutational pathogenicity.

### Mutations at phosphorylation sites within PS IDRs are mostly benign

Protein phosphorylation is used biologically as a mechanism to control many critical life processes (74, 75) and phosphorylation often occurs within protein regions that are ID (76). Phosphorylation has been found to both promote (77, 78) and inhibit (79, 80) phase separation by modulating protein charge, flexibility, and oligomerization state.

We found that 18.8% of human proteins with confirmed phosphorylation sites were predicted by ParSe to exhibit phase separation behavior; a 1.1-fold enrichment relative to the reference human proteome (Figure S5). The normalized rate for confirmed phosphorylation sites found in predicted PS IDRs, nonPS IDRs, and folded regions was 2.73 ± 0.07, 1.77 ± 0.07, and 0.67 ± 0.03, respectively (Figure 2A). Thus, protein phosphorylation was highest in predicted PS ID regions or domains.

However, though phosphorylation sites preferentially map to predicted PS IDRs, we found that missense mutations at such sites were overwhelmingly benign. 54 of 32,665 pathogenic missense mutations (0.17%) in the humsavar index were found both at a confirmed phosphorylation site and in a predicted PS IDR, yielding a normalized incident rate of 0.24 ± 0.03 (Figure 2C). The normalized rate for pathogenic missense mutations at a confirmed phosphorylation site and in a predicted nonPS IDR was 0.23 ± 0.03, and in a predicted folded region was 0.99 ± 0.04. The pathogenic versus benign odds ratio was similarly low, 0.26 ± 0.04 and 0.23 ± 0.03, for phosphorylation sites in PS and nonPS IDRs, respectively, when compared to phosphorylation sites in folded regions, 1.0 ± 0.05 (Figure S3C). Thus, pathogenicity was highest for missense mutations at a phosphorylation site that also occurred in a predicted folded region, and substantially more benign (∼4-fold) for mutations at a phosphorylation site in predicted IDRs, both PS and nonPS. Possibly explaining this result, phosphorylation in folded domains has been observed to occur at buried positions that become transiently accessible, which can directly affect protein structure, dynamics, and biological activity (81). In contrast, phosphorylation in IDRs is often characterized by multisite phosphorylation (82–85) and, accordingly, mutational perturbation of a single site might be less deleterious to function.

### Disease-associated proteins are enriched for phase separation

Tsang et al, using the PScore algorithm (46), found that disease-associated human proteins are enriched in phase separation potential (27). Similarly, ParSe predicts that ∼23% of human proteins annotated as disease-associated in UniProt are likely to exhibit homotypic phase separation behavior, which is a 1.4-fold enrichment compared to the reference human proteome (Figure S5).

As an alternative approach to quantify the differences in PS potential between protein sets, the percent of set data were used to create recall plots with the reference human proteome as the comparison set and from which the area under the curve (AUC) is calculated (Figure S8). This alternative approach was used because, first, when comparing sequence sets, it eliminates the need for a cutoff PS potential for defining proteins as PS or nonPS. The computed AUC is proportional to the fold enrichment for PS whether the cutoff is 25, 50, 100, etc. (Figure S9). AUC > 0.5 indicates a sequence set enriched in computed phase separation potential relative to the reference human proteome. Secondly, the computed AUC can be compared to the mean AUC from sets of random human proteins, facilitating comparisons between sequence sets of vastly different sizes (Figure 3A). AUC values more than two standard deviations from the mean AUC of random sets were statistically equivalent to *p*-values < 0.05 (Figure 3A inset). Thus, AUC values, when compared to the random expectation, can identify when sets of PS potentials have statistically significant differences.

**Figure 3.**
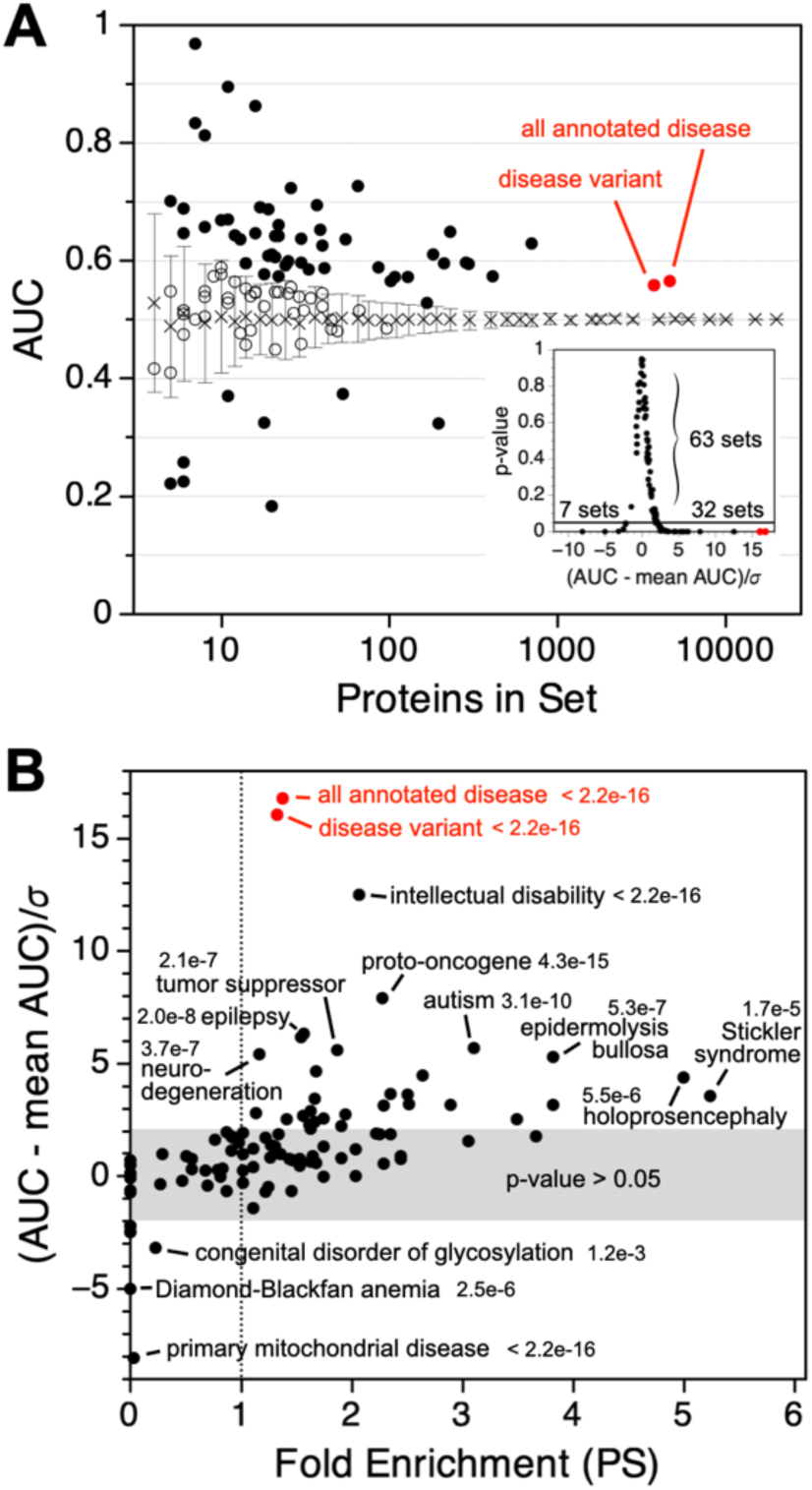
Disease-associated proteins are enriched for phase separation. (**A**) AUC for sets of proteins annotated as disease associated in UniProt were plotted according to the number of proteins in each set (open and filled circles). The mean and standard deviation (x with error bars) from random protein sets were determined by sampling the reference human proteome one hundred times at each indicated set size. Red circles show AUC for the full set of disease-associated proteins and a set containing proteins for which at least one genetic variant involved in a disease has been reported. The inset shows the relationship between the *p*-value and the number of standard deviations from the mean random AUC. Both AUC and the *p*-value were obtained by comparing the distribution of PS potential in a set against its distribution in the reference human proteome. (**B**) Fold enrichment for PS is plotted according to the number of standard deviations from the mean random AUC. For labeled outliers, the *p*-value is shown.

This approach was used to analyze the sets of disease-associated human proteins (Table S3). When sub-grouped by disease type, 79 of 100 disease types produced AUC > 0.5 (Figure S10), 51 produced AUC more than a standard deviation above the mean AUC from random proteins (Figure 3A), and 32 produced AUC more than 2 standard deviations above the mean random AUC (Figure 3A inset). In particular, this result recapitulates the previous finding (27) that proteins associated with cancer and autism spectrum disorder have an over-representation of predicted phase-separating behavior (Figure 3B and Table S3).

We used the standard deviation to identify disease types with unusual and extreme enrichment for PS as compared to random. This was done to avoid sub-groups with enrichment for PS (> 1-fold) but also *p*-values > 0.05, indicating data not sufficient to reject the null hypothesis. Several disease sub-groups had significantly elevated AUC values that were more than three standard deviations above the mean of matched random protein sets. These included neurodevelopmental disorders (such as intellectual disability, autism spectrum disorder, epilepsy, microcephaly, and holoprosencephaly), cancer (proto-oncogenes and tumor suppressors), and connective tissue diseases (such as Ehlers-Danlos syndrome) (Figure 3B and Table S3). Many of the proteins associated with cancers, neurodegeneration, and neurodevelopment disorders are involved in synaptic function (86), transcription regulation (87), and chromatin remodeling (88, 89); biological processes that have been linked to phase separation (90–94). Also, phase separation is thought to drive skin barrier formation (95). Thus, aberrant phase separation of skin proteins potentially has a role in skin disease (epidermolysis bullosa, palmoplantar keratoderma, Ehlers-Danlos syndrome).

In comparison, disease sub-groups with computed AUC three standard deviations (or more) below the mean AUC of similarly sized random protein sets were, in order of statistical significance: primary mitochondrial disease, Diamond-Blackfan anemia, and congenital disorder of glycosylation. In addition to lacking predicted PS IDRs, the proteins in these three sub-groups were on average shorter and more folded, ∼370 residues and ∼85% folded, compared to proteins in the reference proteome, ∼560 residues and ∼75% folded, and especially when compared to human proteins *in vitro* confirmed for phase separation, ∼680 residues and ∼50% folded.

### Sequence dependence to pathogenicity does not depend on region identity

We next sought to determine if pathogenic and benign single amino acid missense mutations exhibit substitution-specific preferences, and whether those preferences varied between the region classes. To achieve this, we compared the frequencies of amino acid specific substitutions for pathogenic versus benign and found that the types of mutations that tended to be pathogenic in predicted PS IDRs also tended to be pathogenic in the other protein regions as well.

We first considered the entire set of mutations in the humsavar index (32,665 pathogenic mutations from 3,345 proteins) and found that the most frequent pathogenic substitutions primarily involved charged residues (R→{C, H, Q, W}, G→R, and E→K) as well as proline (P→L and L→P) (Figure S11A). Many of these same substitutions (e.g., R→H, R→Q, P→L, E→K) also were found in mutations of benign clinical significance (39,656 benign mutations from 11,682 proteins), emphasizing the importance of sequence context (Figure S11B). Figure 4A shows the difference in the calculated substitution frequencies, pathogenic minus benign, for all combinations of substitution types. The highest net positive values (i.e., primarily pathogenic) were the nonconservative changes of L→P, G→R, and R→W, while he lowest net negative values (i.e., primarily benign) were substitutions with generally minimal changes in chemical character (V→I, I→V, A→T, and T→A).

**Figure 4.**
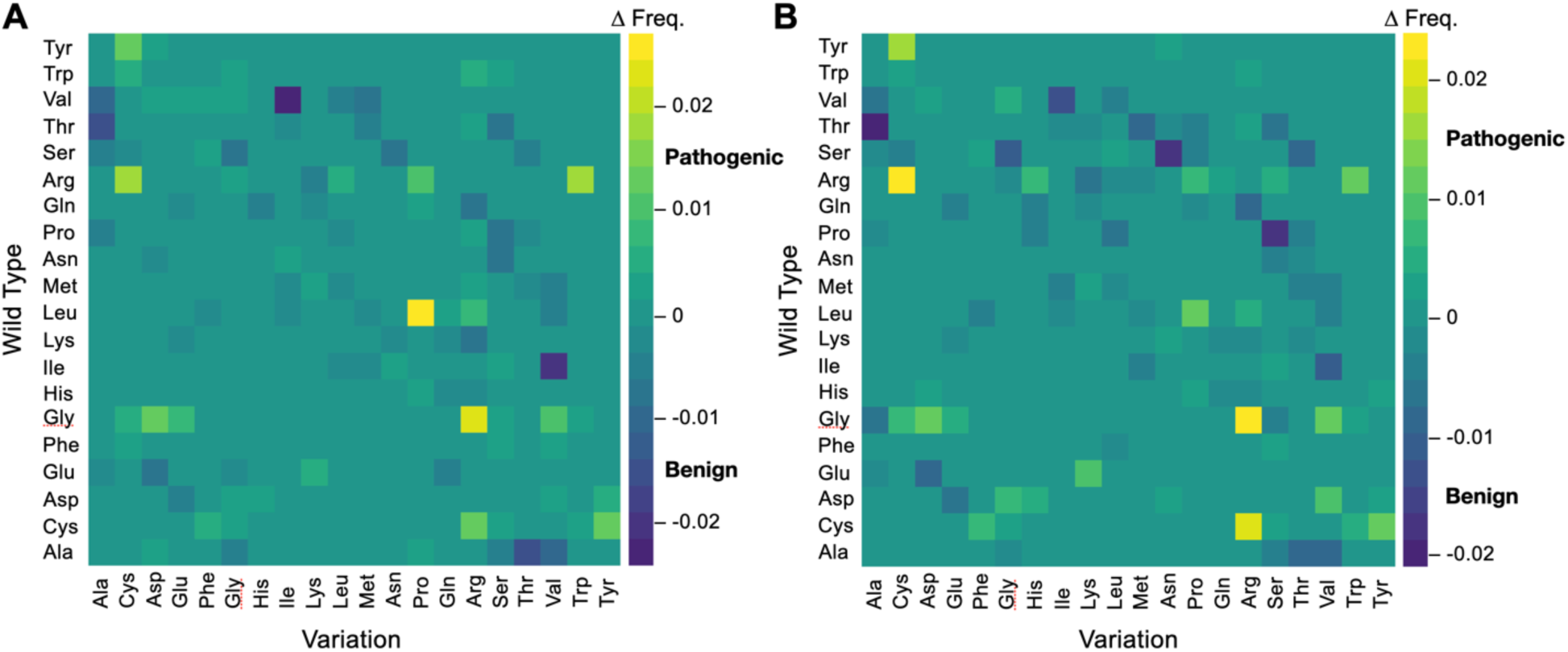
Difference frequencies of pathogenic minus benign single amino acid missense mutations. (**A**) All single amino acid missense mutations in the humsavar index, and (**B**) only those found in a protein region matching the PS ID class and excluding collagen proteins. Difference frequencies (Δ Freq.) were colored according to the scale on the right of each plot.

We next tested if pathogenic and benign substitution trends differed by region identity, to explore why the same substitution sometimes was harmless and sometimes harmful. We found that single amino acid missense mutations in regions identified by ParSe as matching the PS ID class reported different substitution preferences (Figure S12). Here, pathogenic mutations were strongly biased for substitutions from Gly, representing 40% (882 of 2208 substitutions) compared to 11% (496 of 4582 substitutions) of benign mutations. These pathogenic Gly substitutions were primarily from 27 collagen proteins (73%; 642 of 882). Substitutions from Gly would be destabilizing to the collagen triple helix, which structurally requires a repeating Gly-X-Y motif (96, 97). We removed the 27 collagen proteins from our analysis to reduce this bias for substitutions from Gly (removed 4%; 27 of 647 proteins with pathogenic mutations located in predicted PS regions). The resulting difference heat map of pathogenic and benign mutation frequencies (Figure 4B) shows the most net pathogenic changes involved arginine (G→R, R→C, and C→R), while the most net benign involved the small polar hydroxyl-containing residues (T→A, S→A, S→N, and P→S). We used the Mantel test (98) to statistically compare the matrices of net pathogenic and net benign substitutions, for all positions versus PS IDR positions (Figures 4A and 4B), and found a strong, positive correlation coefficient of 0.87 (*p*-value = 0.001), indicating that the patterns of values were highly similar.

Mutations from or to an amino acid type were generally both net pathogenic or both net benign when found in predicted PS IDRs (Figure 5). This was calculated by summing the pathogenic versus benign difference frequencies for each amino acid type, across a row in Figure 4B to obtain the net variation from an amino acid type, and down a column to obtain the net variation to an amino acid type. Exceptions to the overall trend include Gly, which was strongly pathogenic for variations from Gly but not to Gly (even with collagen proteins removed), and Pro, which was often benign for variations from Pro but not to Pro. Single amino acid missense mutations involving Arg and the aromatics, Tyr, Trp, and Phe, were among the net pathogenic group (grey shading in Figure 5). Mutagenesis studies of proteins exhibiting homotypic phase separation behavior have found that Arg and the aromatic types, especially Tyr and Phe, contribute to cohesive protein-protein interactions that drive droplet formation (52–54). These findings are consistent with the idea that mutation at sites implicated in driving phase separation could lead to dysregulation of cellular processes that depend on phase separation, affecting human health. Single amino acid missense mutations involving Ser, Thr, and Ala, in that rank order, were strongly net benign.

**Figure 5.**
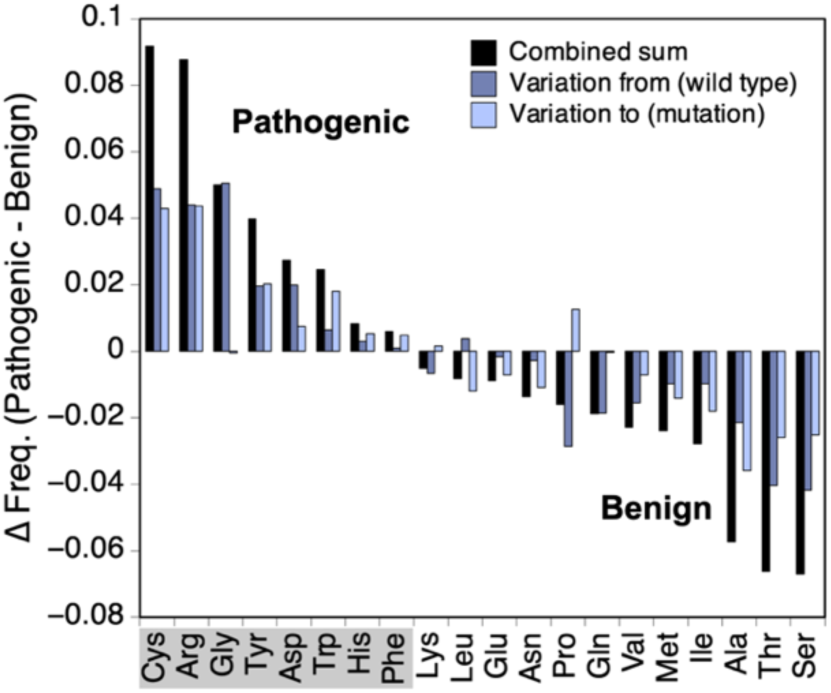
Pathogenicity by amino acid substitution in PS IDRs. Pathogenic minus benign difference frequencies (Δ Freq.) for positions in predicted PS IDRs were summed by wild type amino acid (blue), mutation amino acid (light blue), and the combined sum (black). Contributions from collagen proteins were omitted.

For mutations in regions matching the nonPS ID and folded classes (Figure S13), we again found strong, positive correlations, 0.74 (*p*-value = 0.001) and 0.78 (*p*-value = 0.001), respectively, in the patterns of substitutions that were net pathogenic and net benign when compared to missense mutations found in PS IDRs. This shows that despite robust compositional differences between the three classes of protein regions, PS ID, nonPS ID, and folded (43), the types of mutations that were pathogenic and the types that were benign did not exhibit significant variations by region class. Generally, substitutions involving Arg, Cys, Gly, Tyr, Trp, and Asp were pathogenic, while Ala, Ser, Thr, Val, and Ile were benign.

### Predicting pathogenicity of missense variations by protein region class

We next asked if the pathogenic minus benign difference frequencies could be used to predict clinical significance from the substitution. While more sophisticated predictors of pathogenicity have been developed, typically focused on folded domains (99) or evolutionary models (100, 101), our goal was to leverage our observations of mutations in predicted PS IDRs to extrapolate the significance of the many missense mutations of unknown clinical significance in these regions, in particular.

First, considering all protein regions, we used the difference frequency data in Figure 4A and found that 65% of confirmed pathogenic single amino acid missense mutations were from a substitution type with a net positive difference frequency (i.e., net pathogenic), whereas 63% of confirmed benign single amino acid missense mutations were from a substitution type with a net negative difference frequency (Figure 6A). The combined true positive (correctly predicting pathogenic) and true negative (correctly predicting benign) rate was 64%. Thus, single amino acid missense substitutions were marginally predictive from difference frequency data when applied to the training set, with a 36% error rate. When the substitution trends from the humsavar index were used to predict clinical significance in the natural variance index, the combined true positive and true negative rate was similar, 62% (Figure S14).

**Figure 6.**
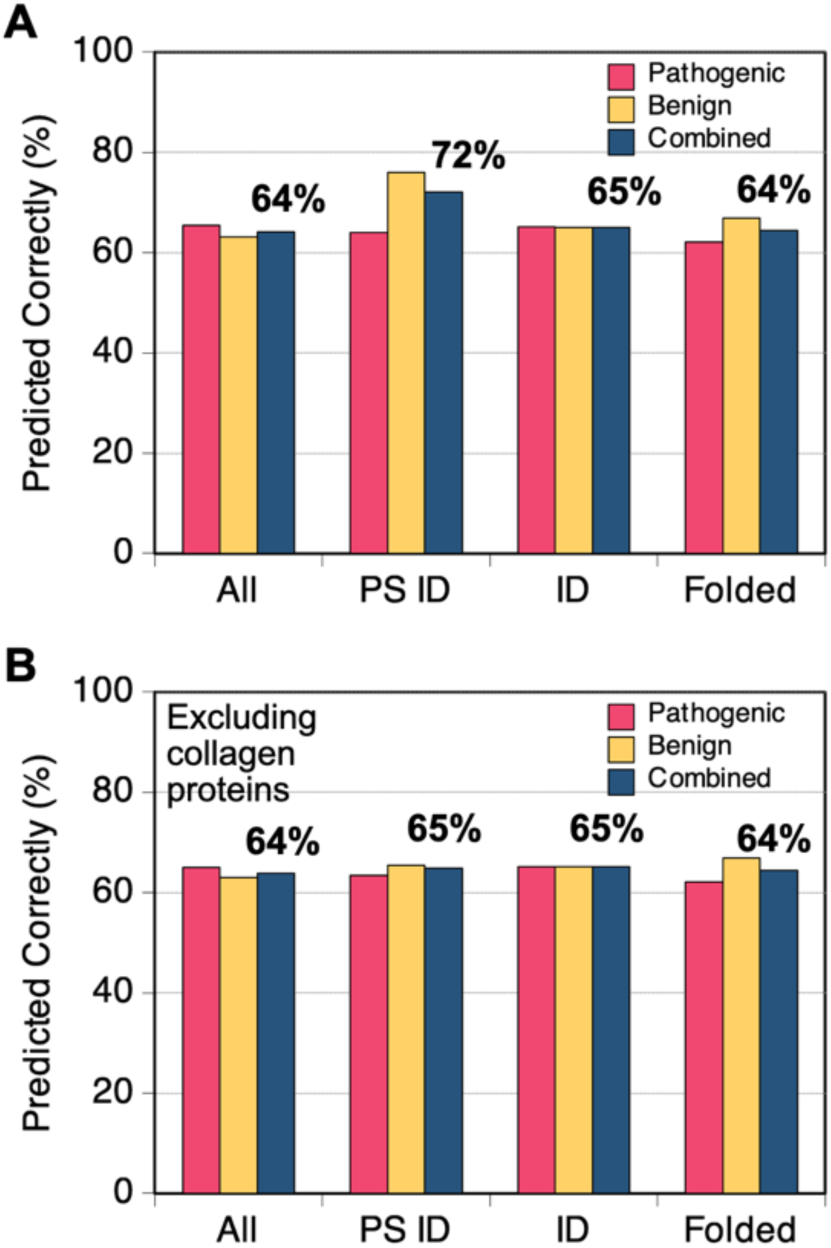
Predicting clinical significance of single amino acid missense mutations. (**A**) Pathogenic minus benign difference frequencies were calculated for all single amino acid missense mutations in the humsavar index and for the subsets residing in protein regions matching the PS ID, ID (i.e., nonPS ID), or folded classes. Clinical significance was predicted for missense mutations with known clinical significance where a positive difference frequency predicts pathogenic, and a negative difference frequency predicts benign. Shown is the percent predicted correctly. (**B**) The effect from excluding collagen proteins.

Focusing on the PS ID class, the combined true positive and true negative rate of missense substitutions was higher, 72% (Figure 6A), owing to a true negative rate that was markedly greater than the true positive rate (76% versus 64%). In protein regions matching the ID and folded classes, the combined true positive and true negative rates were 65% and 64%, respectively. In these calculations, difference frequencies were determined separately for each region class and the substitution-specific difference frequencies for a region class were used to predict clinical significance in that region class. Of note, when the contribution of collagen proteins was omitted from the calculated frequencies and excluded from the prediction tests, the observed predictability in PS IDRs for clinical significance was reduced to the same level found in the other protein region classes (Figure 6B). Thus, though pathogenicity was highest in folded regions for single amino acid missense mutations (Figure 1A), and ∼3-fold higher in PS IDRs relative to nonPS IDRs, the overall predictability of clinical significance from region-specific substitution frequencies was mostly the same across the three region classes. This result is consistent with our finding that, in general, the net pathogenic and net benign substitution types did not change significantly with region identity.

The substitution frequency trends predict that 51% and 41% of single amino acid missense mutations with uncertain clinical significance in the humsavar and natural variants indices, respectively, and 52% and 39% within predicted PS IDRs, are pathogenic when excluding collagen proteins. 40% of the 35,019,669 single amino acid substitutions in the natural variants index with no predicted clinical significance, and 39% within predicted PS IDRs, also were predicted pathogenic. Despite the expected high error rate (∼36%), the net of uncertain and unknown clinical significance missense mutations that were predicted to be pathogenic was substantial (∼40-50%). The high error rate in these predictions clearly suggests that clinical significance depends on additional input variables.

## Discussion

It has been proposed that mutations that alter IDR-induced phase separation may have a higher likelihood of causing disease (27–30). Indeed, the results from several studies indicate a role for aberrant physiological protein phase separation in the progression of some human disorders (31–39). In this work, we identified disease and comparison datasets to map the prevalence of phase separation across a wide range of human diseases, finding that many show an enrichment for phase separation in the proteins associated with them. The diseases most strongly linked to phase separation included phenotypes associated with intellectual disability, cancer, neurodegeneration, and the skin (Figure 3B). We also found that genetic missense variations, regardless of clinical significance, occur at higher rates in proteins predicted to exhibit phase separation behavior, which we attribute to the longer average length and higher ID content of predicted PS proteins. Pathogenic missense mutation rates were highest in the folded regions of proteins (Figure 1B), matching prior reports (9). Benign missense mutation rates did not depend on region identity.

Supporting a link between phase separation and disease, pathogenic missense mutations in predicted IDRs were found more frequently in PS as compared to nonPS regions, by a factor of approximately 3 (Figure 1B). Thus, although IDRs, and even PS IDRs, show low evolutionary conservation of sequence, mutations in these regions can be functionally significant. Pathogenicity was highest for missense mutations in a predicted PS IDR when also in a confirmed SLiM, and significantly higher than for missense mutations in a SLiM and either of the other region classes (Figure 2B). Though highest in PS IDRs, pathogenicity increased for mutations at SLiM positions in each region class (Figure S7), possibly reflecting the conserved sequence patterning of SLiMs (102) and their crucial roles in a wide variety of cellular processes (73).

The significantly higher pathogenicity of SLiMs located in PS IDRs compared to those in folded regions may reflect structural accessibility constraints. SLiMs function as short, modular interaction motifs that typically rely on being solvent-exposed to engage with their recognition partners (103). IDRs offer a flexible and accessible environment where SLiMs can participate in transient interactions. SLiMs embedded within folded domains may be conformationally buried and subsequently less accessible for interaction. This suggests that SLiMs may necessitate disorder for flexibility and functionality, which may explain the increased likelihood of pathogenicity when SLiMs are located within PS IDRs.

The finding that pathogenic mutations were not enriched at phosphorylation sites in PS IDRs (Figure 2C) contrasts with the observed enrichment of pathogenic mutations in SLiMs in the same regions. Phosphorylation in IDRs often involves multisite modification rather than dependence on a single site (82–85). In proteins such as tau, hyperphosphorylation across many residues regulates phase separation behavior (79). In contrast, SLiMs generally function as discrete, site-specific modules for molecular recognition, therefore a mutation in a single SLiM can disrupt binding events and lead to more pronounced pathogenic effects. These structural and functional differences between the two sequence elements may help explain why they differ in their susceptibility to pathogenic outcomes following mutation.

The clinical significance of missense mutations in predicted PS IDRs revealed a dependence on amino acid type that largely mirrored the dependence observed in nonPS and folded regions (Figures 4 and S13). Single amino acid substitutions involving Cys, Arg, Gly, Asp, His, and the aromatics, Tyr, Phe, and Trp, were mostly pathogenic, whereas substitutions involving the other common amino acids were mostly benign (Figure 5). Cys residues are known to promote protein phase separation under oxidizing conditions (104). Numerous mutagenesis studies find Arg, Tyr, and Phe as mediators of droplet formation (46, 51–54). Mutating charge patterns through Asp substitution has been found to modulate phase separation potential (51). Protein-protein interactions involving His have been shown to contribute to phase separation behavior (105), especially toward pH responsive self-assembling processes (106). We conclude that this agreement between amino acid types known to facilitate protein phase separation and the pathogenicity of missense mutations found in predicted PS IDRs reflects the adverse effects of mutating residues that modulate the stability, function, and properties of biological condensates. This work also provides a foundation from which to optimize prediction of the effects of mutations on phase separation, which has historically been difficult.

The computed difference frequencies, pathogenic minus benign, by substitution type for missense mutations in predicted PS IDRs correctly assigned clinical significance to 72% of humsavar missense mutations residing in a predicted PS IDR, and 65% when excluding collagen proteins. From this, we conclude that the properties and functions of IDR-driven biological condensates are tuned by the bulk composition of the corresponding PS IDRs. Efforts to improve prediction of pathogenicity in these domains will likely require additional input variables that provide cellular context, e.g., localization, interacting partners, and gene ontology.

## Methods

### Protein Databases

The one-sequence-per-gene version of the reference human proteome, UP000005640, was downloaded from the Universal Protein Knowledgebase (55), UniProt, in November 2023. This proteome was used as the comparison protein set. Disease-associated human proteins were obtained by searching the UniProt database for proteins annotated as “Human”, “Reviewed (Swiss-Prot)”, and the keyword “Disease”, also in November 2023. The proteins in this set were grouped by disease type by UniProt. Human proteins with confirmed SLiMs were found by search of the Eukaryotic Linear Motif database (61), in January 2024. Human proteins with confirmed phosphorylation sites were obtained from the Eukaryotic Phosphorylation Site Database 2.0 (62) in March 2024. Human proteins with confirmed IDRs were found by search of the DisProt database (56), in June 2025. Human proteins with confirmed phase separation behavior were found by search of the CD-CODE database (57), in June 2025.

### Genetic Variations Databases

The human variants index, file humsavar.txt release 2023_05 of 08-Nov-2023, and the natural variants index, file homo_sapiens_variation.txt release 2023_05 of 11-Oct-2023, both were downloaded from ftp.uniprot.org in November 2023.

### Sequence-calculated Phase Separation Potential

Calculations of phase separation potential for individual protein sequences used the ParSe algorithm, version 2, described elsewhere (43) and available as a webtool (50) at stevewhitten.github.io/Parse_v2_web. Calculations of phase separation potential for sequence sets in the FASTA format, yielding AUC and percent of set values reported in this study, used the webtool at stevewhitten.github.io/ParSe_v2_FASTA_reviewed_proteome with source code available at github.com/stevewhitten.

### Mann-Whitney *U*-test

One-tail *p*-values were calculated using the Mann-Whitney *U*-test (69) as implemented in the R statistical computing and graphics program (107).

### Mantel test

Mantel statistics (98) for comparing matrices were calculated by Pearson’s product-moment correlation and with 999 permutations. These calculations used the vegan package (108) as implemented in the R statistical computing and graphics program (107).

### Fold Enrichment

Fold Enrichment for a protein set was calculated as the fractional number of PS predicted proteins in a set divided by the fractional number of PS predicted proteins in the reference proteome. Protein type, PS or nonPS, was determined by the ParSe algorithm, version 2, where PS proteins had PS potential ≥100.

### Calculation of Normalized Missense Mutation Rate

The rate represents a proportion, *p*, of mutations found in a region,

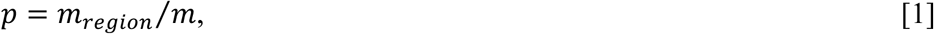

where *m* is the total number of mutations, *m_region_* is the subset of *m* found in a region, and regions can be PS IDRs, nonPS IDRs, folded regions, SLiMs in PS IDRs, etc. Pathogenic rates were calculated using missense mutations labeled likely pathogenic or pathogenic; benign rates used missense mutations labeled likely benign or benign. Next, *p* was multiplied by a factor, *k*, that normalized rates according to the proportion of residues found in a region. Specifically,

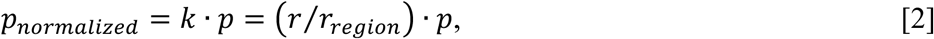

where *r* is the total number of residues in the sequence set (e.g., the reference proteome), and *r_region_* is the subset of *r* found in a region type. The standard error of the proportion (non-scaled) is,

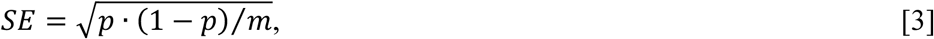

The adjusted standard error for normalized *p* is,

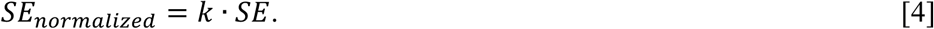

### Calculation of SLiM and Phosphorylation Rate

Rates of SLiMs and phosphorylation by region were calculated in a manner identical to the mutation rate. For SLiMs, equation 1 was modified as,

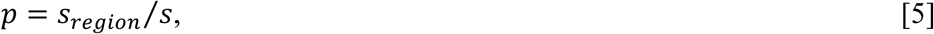

where *s* is the total number of SLiMs and *s_region_* the subset found in a region, and for phosphorylation as,

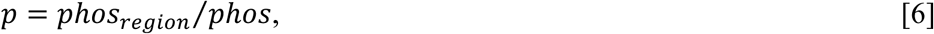

where *phos* is the total number of phosphorylation sites and *phos_region_* the subset found in a region. Each rate was normalized by *k* using equation 2. For SLiMs, SE was calculated using equation 3 with *s* substituted for *m*. For phosphorylation, SE was calculated with *phos* substituted for *m*. The adjusted standard error was calculated as given by equation 4.

### Calculation of Pathogenic versus Benign Odds Ratio

Odds ratios were used to compare the probabilities of pathogenic versus benign mutations. Probabilities were determined by,

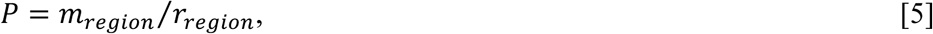

were *m_region_* and *r_region_* are defined above. Pathogenic probabilities were calculated using missense mutations labeled likely pathogenic or pathogenic; benign probabilities used missense mutations labeled likely benign or benign. The odds ratio, *OR*, was determined by,

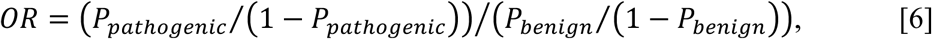

The standard error for the natural log of *OR* is (109),

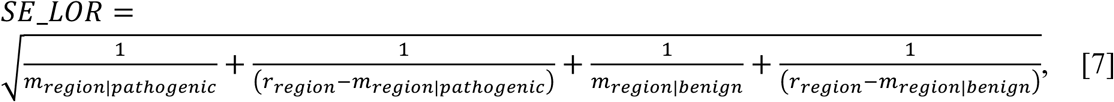

where *m_region|pathogenic_* and *m_region|benign_* represent the number of pathogenic and benign mutations found in a region, respectively. The standard error for the OR was estimated by (110),

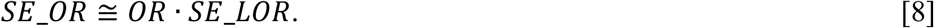

### Calculation of Substitution Frequencies

Substitution frequencies were calculated as the fraction of single amino acid missense mutations of a given type, e.g., Gly-to-Ala, in a variant index,

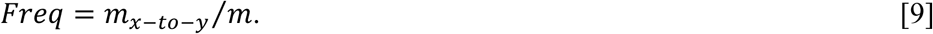

Substitution frequencies by region class, for PS ID, nonPS ID, and folded, were calculated as the fraction of single amino acid missense mutations of a given type, e.g., Gly-to-Ala, in a variant index for those found in a region class,

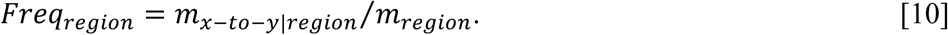

Protein region class by position in a primary sequence was determined using ParSe version 2.

## Data Availability

Source code for both the ParSe v2 algorithm and its webtool applications can be downloaded from github.com/stevewhitten.

## Additional Information

### Supporting Information

This article contains supporting information (43, 55, 61, 62, 69, 111, 112).

### Author Contributions

L. E. H. and S. T. W. conceptualization; O. L. K., K. A. L., L. E. H., and S. T. W. formal analysis; O. L. K., K. A. L., L. E. H., and S. T. W. investigation; L. E. H. and S. T. W. methodology; S. T. W. writing–original draft; O. L. K., K. A. L., L. E. H., and S. T. W. writing–review and editing.

### Abbreviations

The abbreviations used are:

AUC: area under the curve
ELM: Eukaryotic Linear Motif
ID: intrinsically disordered
IDR: intrinsically disordered region
nonPS: nonphase-separating
PS: phase-separating
SLiM: short linear motif
UniProt: Universal Protein Knowledgebase.

