## supporting information for "Missense mutations in intrinsically disordered protein regions link pathogenicity and phase separation"

Short title: Missense mutations in phase-separating IDRs

*Oliver L. Kipp, Karen A. Lewis, Loren E. Hough, and Steven T. Whitten*

### Contents:

Supporting Tables
S1. Human proteome, missense mutation, SLiM, and phosphorylation statistics.
S2. Normalized missense mutation rate calculations.
S3. AUC and *p*-value for protein sets annotated as disease associated in UniProt.

Supporting Figures
S1. The ParSe algorithm (version 2).
S2. Predicted PS potential in the reference human proteome and homotypic PS proteins.
S3. Enrichment for phase separation in protein sets.
S4. Protein length and percent of residues folded in PS and nonPS proteins.
S5. Pathogenic versus benign odds ratios for missense mutations.
S6. Pathogenic missense mutation rate by region class for the natural variants index.
S7. Pathogenic missense mutation rate increases in SLiMs by region class.
S8. Quantifying differences in predicted PS potential between protein sets.
S9. AUC trends with enrichment for phase separation behavior.
S10. AUC rank order for protein sets annotated as disease associated.
S11. Frequency of single amino acid missense variation types.
S12. Frequency of single amino acid missense variation types in PS IDRs.
S13. Pathogenic minus benign difference frequencies for single amino acid missense variation types in nonPS and folded regions.
S14. Predicting clinical significance of missense mutations in the natural variants index.

Supporting References

### Supporting Tables

#### Table S1. Human proteome, missense mutation, SLiM, and phosphorylation statistics.

| | |
|---|---|
| proteins [a] | 20,435 |
| proteins w/ $N \geq 25$ | 20,383 |
| PS proteins w/ $N \geq 25$ | 3,338 |
| nonPS proteins w/ $N \geq 25$ | 17,045 |
| residues | 11,403,806 |
| residues in PS proteins | 3,810,609 |
| residues in nonPS proteins | 7,593,197 |
| residues in PS IDRs | 624,550 |
| residues in nonPS IDRs | 1,061,254 |
| residues in folded regions | 8,378,569 |
| pathogenic mutations [b] | 32,665 |
| pathogenic mutations in PS proteins | 9,083 |
| pathogenic mutations in nonPS proteins | 23,582 |
| pathogenic mutations in PS IDRs | 1,186 |
| pathogenic mutations in nonPS IDRs | 727 |
| pathogenic mutations in folded regions | 29,225 |
| proteins with pathogenic mutations | 3,345 |
| PS proteins with pathogenic mutations | 706 |
| nonPS proteins with pathogenic mutations | 2,639 |
| benign mutations | 39,654 |
| benign mutations in PS proteins | 10,995 |
| benign mutations in nonPS proteins | 28,659 |
| benign mutations in PS IDRs | 2,191 |
| benign mutations in nonPS IDRs | 3,828 |
| benign mutations in folded regions | 28,812 |
| proteins with benign mutations | 11,681 |
| PS proteins with benign mutations | 2,301 |
| nonPS proteins with benign mutations | 9,380 |
| uncertain clinical significance mutations | 10,177 |
| uncertain clinical significance mutations in PS proteins | 3,361 |
| uncertain clinical significance mutations in nonPS proteins | 6,816 |
| uncertain clinical significance mutations in PS IDRs | 335 |
| uncertain clinical significance mutations in nonPS IDRs | 780 |
| uncertain clinical significance mutations in folded regions | 8,149 |
| proteins with uncertain clinical significance mutations | 2,802 |
| PS proteins with uncertain clinical significance mutations | 743 |
| nonPS proteins with uncertain clinical significance mutations | 2,059 |
| SLiMs [c] | 2,247 |
| proteins with a SLiM | 1,370 |
| PS proteins with a SLiM | 523 |
| nonPS proteins with a SLiM | 847 |
| SLiM residues | 16,224 |
| SLiMs in PS proteins | 964 |
| SLiMs in nonPS proteins | 1,282 |
| SLiMs in PS IDRs | 222 |
| SLiM residues in PS IDRs | 1,410 |

| | |
|---|---|
| SLiMs in nonPS IDRs | 533 |
| SLiM residues in nonPS IDRs | 3,668 |
| SLiMs in folded regions | 692 |
| SLiM residues in folded regions | 5,834 |
| pathogenic mutations in SLiMs | 141 |
| pathogenic mutations in SLiMs in PS IDRs | 44 |
| pathogenic mutations in SLiMs in nonPS IDRs | 18 |
| pathogenic mutations in SLiMs in folded regions | 52 |
| benign mutations in SLiMs | 76 |
| benign mutations in SLiMs in PS IDRs | 8 |
| benign mutations in SLiMs in nonPS IDRs | 11 |
| benign mutations in SLiMs in folded regions | 35 |
| uncertain clinical significance mutations in SLiMs | 194 |
| uncertain clinical significance mutations in SLiMs in PS IDRs | 23 |
| uncertain clinical significance mutations in SLiMs in nonPS IDRs | 31 |
| uncertain clinical significance mutations in SLiMs in folded regions | 32 |
| phosphorylation sites [d] | 531,023 |
| proteins with a phosphorylation site | 30,600 |
| PS proteins with a phosphorylation site | 5,763 |
| nonPS proteins with a phosphorylation site | 24,837 |
| phosphorylation sites in PS proteins | 212,256 |
| phosphorylation sites in nonPS proteins | 318,767 |
| phosphorylation sites in PS IDRs | 79,280 |
| phosphorylation sites in nonPS IDRs | 87,645 |
| phosphorylation sites in folded regions | 263,030 |
| pathogenic mutations at phosphorylation sites | 942 |
| pathogenic mutations at phosphorylation sites in PS IDRs | 54 |
| pathogenic mutations at phosphorylation sites in nonPS IDRs | 58 |
| pathogenic mutations at phosphorylation sites in folded regions | 746 |
| benign mutations at phosphorylation sites | 1,542 |
| benign mutations at phosphorylation sites in PS IDRs | 210 |
| benign mutations at phosphorylation sites in nonPS IDRs | 256 |
| benign mutations at phosphorylation sites in folded regions | 746 |
| uncertain clinical significance mutations at phosphorylation sites | 403 |
| uncertain clinical significance mutations at phosphorylation sites in PS IDRs | 45 |
| uncertain clinical significance mutations at phosphorylation sites in nonPS IDRs | 51 |
| uncertain clinical significance mutations at phosphorylation sites in folded regions | 239 |

[a] Human protein sequences were obtained from the UniProt reference human proteome (1). For a protein, ParSe analysis (2) requires a minimum sequence length, $N$, of 25 residues. The table statistics, other than the first row, thus includes contributions only from proteins with $N \geq 25$.
[b] Mutant sites represent the single amino acid missense mutations in the humsavar index (1).
[c] SLiMs sites in human proteins were obtained from the Eukaryotic Linear Motif resource (3).
[d] Phosphorylation sites in human proteins were obtained from the Eukaryotic Phosphorylation Site Database 2.0 (4).

**Table S2. Normalized missense mutation rate calculations.** Input values from Table S1.

pathogenic mutations in folded regions = (29,225/32,665)/(8,378,569/11,403,806) = 1.218
pathogenic mutations in PS IDRs = (1,186/32,665)/(624,550/11,403,806) = 0.663
pathogenic mutations in nonPS IDRs = (727/32,665)/(1,061,254/11,403,806) = 0.239

benign mutations in folded regions = (28,812/39,654)/(8,378,569/11,403,806) = 0.989
benign mutations in PS IDRs = (2,191/39,654)/(624,550/11,403,806) = 1.009
benign mutations in nonPS IDRs = (3,828/39,654)/(1,061,254/11,403,806) = 1.037

SLiMs in folded regions = (692/2,247)/(8,378,569/11,403,806) = 0.419
SLiMs in PS IDRs = (222/2,247)/(624,550/11,403,806) = 1.804
SLiMs in nonPS IDRs = (533/2,247)/(1,061,254/11,403,806) = 2.549

phosphorylation sites in folded regions = (263,030/531,023)/(8,378,569/11,403,806) = 0.674
phosphorylation sites in PS IDRs = (79,280/531,023)/(624,550/11,403,806) = 2.726
phosphorylation sites in nonPS IDRs = (87,645/531,023)/(1,061,254/11,403,806) = 1.774

pathogenic mutations in SLiMs = (141/32,665)/(16,224/11,403,806) = 3.034
pathogenic mutations in SLiMs in folded regions = (52/32,665)/(5,834/11,403,806) = 3.112
pathogenic mutants in SLiMs in PS IDRs = (44/32,665)/(1,410/11,403,806) = 10.894
pathogenic mutants in SLiMs in nonPS IDRs = (18/32,665)/(3,668/11,403,806) = 1.713

benign mutations in SLiMs = (76/39,654)/(16,224/11,403,806) = 1.347
benign mutations in SLiMs in folded regions = (35/39,654)/(5,834/11,403,806) = 1.725
benign mutations in SLiMs in PS IDRs = (8/39,654)/(1,410/11,403,806) = 1.632
benign mutations in SLiMs in nonPS IDRs = (11/39,654)/(3,668/11,403,806) = 0.862

pathogenic mutations at phosphorylation sites = (942/32,665)/(531,023/11,403,806) = 0.619
pathogenic mutations at phosphorylation sites in folded regions = (746/32,665)/(263,030/11,403,806) = 0.990
pathogenic mutations at phosphorylation sites in PS IDRs = (54/32,665)/(79,280/11,403,806) = 0.238
pathogenic mutations at phosphorylation sites in nonPS IDRs = (58/32,665)/(87,645/11,403,806) = 0.231

benign mutations at phosphorylation sites = (1,542/39,654)/(531,023/11,403,806) = 0.835
benign mutations at phosphorylation sites in folded regions = (746/39,654)/(263,030/11,403,806) = 0.816
benign mutations at phosphorylation sites in PS IDRs = (210/39,654)/(79,280/11,403,806) = 0.762
benign mutations at phosphorylation sites in nonPS IDRs = (256/39,654)/(87,645/11,403,806) = 0.840

**Table S3. AUC and *p*-value for protein sets annotated as disease associated in UniProt.**

| Set[a] | Proteins[b] | PS proteins[c] | AUC[d] | mean AUC[e] | $\sigma$[f] | (AUC - mean AUC)/$\sigma$ | *p*-value[g] |
|---|---|---|---|---|---|---|---|
| All | 4638 | 1046 | 0.5627 | 0.5004 | 0.0037 | 16.7677 | < 2.2E-16 |
| Disease variant | 3756 | 817 | 0.5575 | 0.4994 | 0.0036 | 16.0386 | < 2.2E-16 |
| Intellectual disability | 703 | 238 | 0.6286 | 0.4986 | 0.0104 | 12.4926 | < 2.2E-16 |
| Proto-oncogene | 231 | 86 | 0.6482 | 0.4997 | 0.0188 | 7.9015 | 4.31E-15 |
| Deafness | 285 | 73 | 0.5961 | 0.4981 | 0.0156 | 6.2994 | 1.75E-08 |
| Epilepsy | 297 | 75 | 0.5939 | 0.4981 | 0.0156 | 6.1580 | 1.98E-08 |
| Autism spectrum disorder | 65 | 33 | 0.7268 | 0.5011 | 0.0397 | 5.6793 | 3.06E-10 |
| Tumor suppressor | 183 | 56 | 0.6103 | 0.5015 | 0.0194 | 5.5967 | 2.14E-07 |
| Neurodegeneration | 413 | 79 | 0.5725 | 0.4989 | 0.0136 | 5.3997 | 3.73E-07 |
| Epidermolysis bullosa | 16 | 10 | 0.8623 | 0.4904 | 0.0704 | 5.2796 | 5.27E-07 |
| Dwarfism | 211 | 58 | 0.5948 | 0.5013 | 0.0201 | 4.6403 | 1.14E-06 |
| Primary microcephaly | 37 | 16 | 0.6937 | 0.4958 | 0.0444 | 4.4625 | 4.33E-05 |
| Holoprosencephaly | 11 | 9 | 0.8949 | 0.5002 | 0.0905 | 4.3617 | 5.50E-06 |
| Craniosynostosis | 26 | 10 | 0.7226 | 0.5031 | 0.0601 | 3.6542 | 8.55E-05 |
| Palmoplantar keratoderma | 39 | 16 | 0.6514 | 0.4978 | 0.0426 | 3.6020 | 0.0008523 |
| Stickler syndrome | 7 | 6 | 0.9681 | 0.5048 | 0.1304 | 3.5529 | 1.73E-05 |
| Ectodermal dysplasia | 55 | 15 | 0.6356 | 0.4969 | 0.0404 | 3.4356 | 0.0004134 |
| Ehlers-Danlos syndrome | 17 | 7 | 0.6907 | 0.5000 | 0.0597 | 3.1951 | 0.006468 |
| Atrial septal defect | 8 | 5 | 0.8126 | 0.4936 | 0.1008 | 3.1646 | 0.002136 |
| Congenital hypothyroidism | 19 | 9 | 0.6873 | 0.4949 | 0.0610 | 3.1548 | 0.003914 |
| Hypotrichosis | 40 | 15 | 0.6248 | 0.4977 | 0.0408 | 3.1126 | 0.005988 |
| Cone-rod dystrophy | 30 | 8 | 0.6366 | 0.4930 | 0.0496 | 2.8928 | 0.009102 |
| Neuropathy | 129 | 24 | 0.5716 | 0.4994 | 0.0260 | 2.7765 | 0.004794 |
| Premature ovarian failure | 22 | 7 | 0.6604 | 0.5026 | 0.0581 | 2.7178 | 0.008063 |
| Diabetes mellitus | 86 | 22 | 0.5879 | 0.4998 | 0.0331 | 2.6621 | 0.004732 |
| Corneal dystrophy | 21 | 6 | 0.6409 | 0.4883 | 0.0602 | 2.5370 | 0.02663 |
| Alport syndrome | 7 | 4 | 0.8337 | 0.5048 | 0.1304 | 2.5222 | 0.002096 |
| Retinitis pigmentosa | 108 | 25 | 0.5720 | 0.4985 | 0.0293 | 2.5094 | 0.007915 |
| Nephronophthisis | 22 | 6 | 0.6416 | 0.5026 | 0.0581 | 2.3941 | 0.02336 |
| Cardiomyopathy | 102 | 27 | 0.5652 | 0.5008 | 0.0282 | 2.2847 | 0.02953 |
| Age-related macular degeneration | 16 | 5 | 0.6463 | 0.4904 | 0.0704 | 2.2130 | 0.04373 |
| Microphthalmia | 30 | 8 | 0.5963 | 0.4930 | 0.0496 | 2.0810 | 0.05867 |

| | | | | | | | |
|---|---|---|---|---|---|---|---|
| Kallmann syndrome | 21 | 3 | 0.6049 | 0.4883 | 0.0602 | 1.9387 | 0.09627 |
| Oncogene | 12 | 2 | 0.6426 | 0.4965 | 0.0760 | 1.9229 | 0.08336 |
| Thrombophilia | 11 | 4 | 0.6696 | 0.5002 | 0.0905 | 1.8718 | 0.04845 |
| Parkinsonism | 41 | 9 | 0.5863 | 0.4987 | 0.0470 | 1.8634 | 0.05735 |
| Schizophrenia | 19 | 7 | 0.6080 | 0.4949 | 0.0610 | 1.8546 | 0.106 |
| Myofibrillar myopathy | 13 | 5 | 0.6355 | 0.4928 | 0.0777 | 1.8375 | 0.09324 |
| Peters anomaly | 5 | 3 | 0.7007 | 0.4879 | 0.1202 | 1.7706 | 0.1244 |
| SCID | 20 | 3 | 0.6099 | 0.4989 | 0.0629 | 1.7634 | 0.08259 |
| Amelogenesis imperfecta | 25 | 4 | 0.5984 | 0.5071 | 0.0534 | 1.7109 | 0.08422 |
| Hemolytic uremic syndrome | 10 | 2 | 0.6684 | 0.5049 | 0.0956 | 1.7095 | 0.06941 |
| Hypogonadotropic hypogonadism | 33 | 5 | 0.5841 | 0.4931 | 0.0538 | 1.6909 | 0.09757 |
| Hereditary nonpolyposis colorectal cancer | 8 | 1 | 0.6570 | 0.4936 | 0.1008 | 1.6212 | 0.1243 |
| Emery-Dreifuss muscular dystrophy | 6 | 3 | 0.6880 | 0.5098 | 0.1144 | 1.5573 | 0.1042 |
| Ciliopathy | 168 | 27 | 0.5279 | 0.4997 | 0.0192 | 1.4734 | 0.229 |
| Parkinson disease | 24 | 5 | 0.5915 | 0.4992 | 0.0670 | 1.3778 | 0.1168 |
| Congenital stationary night blindness | 14 | 4 | 0.5951 | 0.5040 | 0.0695 | 1.3107 | 0.1922 |
| Glaucoma | 14 | 3 | 0.5946 | 0.5040 | 0.0695 | 1.3035 | 0.2039 |
| Systemic lupus erythematosus | 22 | 4 | 0.5729 | 0.5026 | 0.0581 | 1.2112 | 0.2298 |
| Cockayne syndrome | 6 | 2 | 0.6462 | 0.5098 | 0.1144 | 1.1920 | 0.2089 |
| Hereditary hemolytic anemia | 40 | 6 | 0.5434 | 0.4977 | 0.0408 | 1.1198 | 0.328 |
| Xeroderma pigmentosum | 9 | 2 | 0.5729 | 0.4886 | 0.0858 | 0.9832 | 0.3938 |
| Alzheimer disease | 18 | 3 | 0.5768 | 0.5056 | 0.0728 | 0.9775 | 0.256 |
| Albinism | 21 | 1 | 0.5461 | 0.4883 | 0.0602 | 0.9614 | 0.4661 |
| Asthma | 12 | 1 | 0.5635 | 0.4965 | 0.0760 | 0.8820 | 0.4253 |
| Hirschsprung disease | 10 | 4 | 0.5876 | 0.5049 | 0.0956 | 0.8646 | 0.379 |
| Osteogenesis imperfecta | 26 | 7 | 0.5549 | 0.5031 | 0.0601 | 0.8629 | 0.2869 |
| Williams-Beuren syndrome | 29 | 6 | 0.5384 | 0.4926 | 0.0559 | 0.8206 | 0.4594 |
| Long QT syndrome | 16 | 5 | 0.5468 | 0.4904 | 0.0704 | 0.8004 | 0.4959 |
| Aortic aneurysm | 16 | 4 | 0.5445 | 0.4904 | 0.0704 | 0.7677 | 0.5055 |
| Glycogen storage disease | 22 | 2 | 0.5468 | 0.5026 | 0.0581 | 0.7618 | 0.4351 |
| Lissencephaly | 34 | 8 | 0.5358 | 0.5004 | 0.0475 | 0.7435 | 0.4132 |

| | | | | | | | |
|---|---|---|---|---|---|---|---|
| Brugada syndrome | 10 | 4 | 0.5759 | 0.5049 | 0.0956 | 0.7423 | 0.3917 |
| Mucopolysaccharidosis | 14 | 0 | 0.5521 | 0.5040 | 0.0695 | 0.6917 | 0.507 |
| Congenital myasthenic syndrome | 25 | 6 | 0.5435 | 0.5071 | 0.0534 | 0.6822 | 0.4091 |
| Atrial fibrillation | 15 | 4 | 0.5396 | 0.4885 | 0.0794 | 0.6437 | 0.5408 |
| Limb-girdle muscular dystrophy | 40 | 11 | 0.5214 | 0.4977 | 0.0408 | 0.5812 | 0.6741 |
| Atherosclerosis | 8 | 3 | 0.5478 | 0.4936 | 0.1008 | 0.5381 | 0.6379 |
| Glutaricaciduria | 5 | 0 | 0.5470 | 0.4879 | 0.1202 | 0.4916 | 0.7078 |
| Primary hypomagnesemia | 8 | 2 | 0.5384 | 0.4936 | 0.1008 | 0.4448 | 0.7366 |
| Osteopetrosis | 11 | 2 | 0.5362 | 0.5002 | 0.0905 | 0.3976 | 0.6242 |
| Obesity | 66 | 9 | 0.5144 | 0.5011 | 0.0397 | 0.3346 | 0.7245 |
| Cushing syndrome | 11 | 1 | 0.5280 | 0.5002 | 0.0905 | 0.3069 | 0.6752 |
| Amyloidosis | 31 | 4 | 0.5148 | 0.4997 | 0.0538 | 0.2813 | 0.8068 |
| Leber congenital amaurosis | 27 | 3 | 0.5102 | 0.4980 | 0.0503 | 0.2421 | 0.8552 |
| Intrahepatic cholestasis | 18 | 3 | 0.5221 | 0.5056 | 0.0728 | 0.2262 | 0.6838 |
| Aicardi-Goutieres syndrome | 8 | 0 | 0.5044 | 0.4936 | 0.1008 | 0.1076 | 0.9445 |
| Bartter syndrome | 6 | 0 | 0.5149 | 0.5098 | 0.1144 | 0.0446 | 0.9116 |
| Congenital erythrocytosis | 6 | 2 | 0.5094 | 0.5098 | 0.1144 | -0.0035 | 0.9408 |
| Dystonia | 45 | 6 | 0.4985 | 0.4996 | 0.0451 | -0.0235 | 0.9486 |
| Pseudohermaphroditism | 7 | 2 | 0.4998 | 0.5048 | 0.1304 | -0.0382 | 0.9221 |
| Peroxisome biogenesis disorder | 15 | 0 | 0.4806 | 0.4885 | 0.0794 | -0.0990 | 0.8703 |
| Dyskeratosis congenita | 13 | 1 | 0.4760 | 0.4928 | 0.0777 | -0.2165 | 0.8201 |
| Allergen | 6 | 1 | 0.4739 | 0.5098 | 0.1144 | -0.3137 | 0.7693 |
| Leukodystrophy | 45 | 2 | 0.4834 | 0.4996 | 0.0451 | -0.3583 | 0.7107 |
| Cataract | 96 | 11 | 0.4842 | 0.4964 | 0.0293 | -0.4176 | 0.668 |
| Ichthyosis | 49 | 10 | 0.4802 | 0.4997 | 0.0398 | -0.4912 | 0.8082 |
| Gangliosidosis | 5 | 0 | 0.4095 | 0.4879 | 0.1202 | -0.6525 | 0.4818 |
| Fanconi anemia | 21 | 5 | 0.4487 | 0.4883 | 0.0602 | -0.6574 | 0.5252 |
| Heterotaxy | 14 | 2 | 0.4567 | 0.5040 | 0.0695 | -0.6815 | 0.6306 |
| Congenital muscular dystrophy | 30 | 6 | 0.4582 | 0.4930 | 0.0496 | -0.7009 | 0.4323 |
| Diabetes insipidus | 4 | 0 | 0.4158 | 0.5281 | 0.1514 | -0.7415 | 0.5775 |
| Nemaline myopathy | 11 | 2 | 0.3698 | 0.5002 | 0.0905 | -1.4414 | 0.1357 |
| Congenital adrenal hyperplasia | 6 | 0 | 0.2572 | 0.5098 | 0.1144 | -2.2074 | 0.04577 |

| | | | | | | | |
|---|---|---|---|---|---|---|---|
| Congenital generalized lipodystrophy | 5 | 0 | 0.2214 | 0.4879 | 0.1202 | -2.2178 | 0.04105 |
| Dystroglycanopathy | 18 | 0 | 0.3246 | 0.5056 | 0.0728 | -2.4867 | 0.009927 |
| Chronic granulomatous disease | 6 | 0 | 0.2246 | 0.5098 | 0.1144 | -2.4923 | 0.01442 |
| Congenital disorder of glycosylation | 53 | 2 | 0.3737 | 0.5014 | 0.0401 | -3.1869 | 0.001243 |
| Diamond-Blackfan anemia | 20 | 0 | 0.1824 | 0.4989 | 0.0629 | -5.0293 | 2.49E-06 |
| Primary mitochondrial disease | 197 | 1 | 0.3227 | 0.4982 | 0.0217 | -8.0974 | < 2.2E-16 |

[a] Disease-associated human proteins were annotated and grouped by UniProt (1). "All" refers to the full set of disease-associated proteins and "Disease variant" is the set containing all proteins for which at least one genetic variant involved in a disease has been reported.
[b] Number of proteins in the set with sequence length at least 25 residues, the minimum sequence length required for ParSe analysis (2).
[c] Number of ParSe-predicted PS proteins in the set.
[d] AUC for the set using the reference human proteome as the comparison set.
[e] AUC mean from one hundred random human protein sets using the set size given in column 2.
[f] AUC standard deviation from one hundred random human protein sets using the set size given in column 2.
[g] One-tail *p*-value determined by the Mann-Whitney *U*-test (5) for the calculated PS potentials of the set and using the reference human proteome for the comparison set.

### Supporting Figures

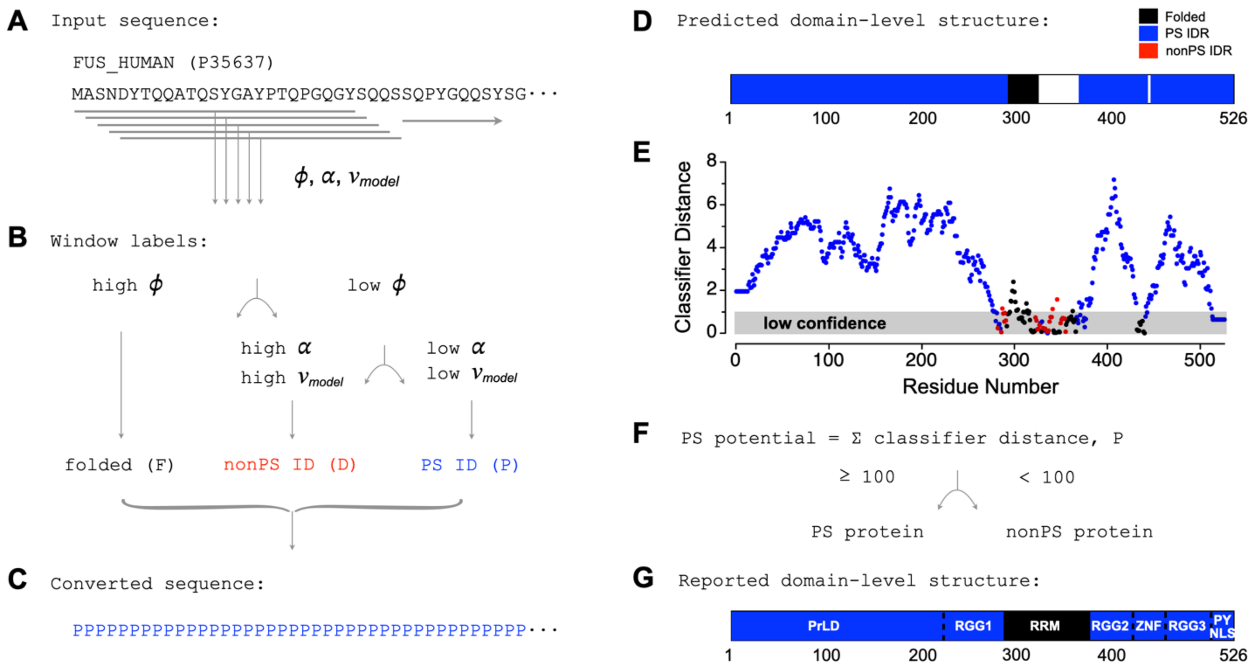

**Figure S1. The ParSe algorithm (version 2).** (**A**) Hydrophobicity ($\phi$), α-helix propensity ($\alpha$), and $\nu_{model}$ are calculated for each contiguous stretch of 25-residues, or "window", in the primary sequence. (**B**) Each window is assigned a label, F, D, or P, depending on the values of $\phi$, $\alpha$, and $\nu_{model}$. (**C**) Window labels are assigned to the central residue of the window. Terminal residues are assigned labels according to the first and last windows. (**D**) Contiguous regions of at least 20 residues that are 90% of only one label P, D, or F are colored blue, red, or black, respectively, to represent predicted PS IDRs, IDRs, or folded regions. White corresponds to regions with mixed P, D, or F labels. (**E**) Classifier distance of each window, assigned to the central residue, and colored according to its label P (blue), D (red), or F (black). This "distance" represents standard deviations from the high/low cutoff values of $\phi$, $\alpha$, and $\nu_{model}$, where standard deviations were determined from the distributions of $\phi$, $\alpha$, and $\nu_{model}$ in the sequences that trained ParSe v2. Low values (<1) indicate low statistical confidence for a window label. (**F**) The classifier distance sum from P-labeled windows is used as a "PS potential"; primary sequences with values ≥100 are PS predicted proteins. (**G**) Reported domain-level organization for the FUS RNA-binding protein (6, 7).

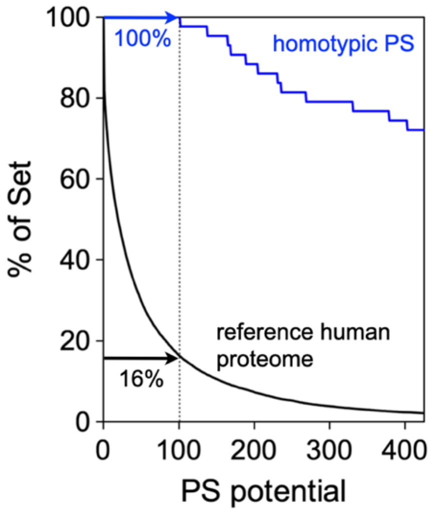

**Figure S2. Predicted PS potential in the reference human proteome and homotypic PS proteins.** The percent of proteins in a set with sequence-calculated PS potential equal to or greater than the value indicated by the x-axis is shown for confirmed homotypic phase-separating (PS) proteins (blue) and the reference human proteome (black). The percent of each set with PS potential ≥100 is indicated in the figure.

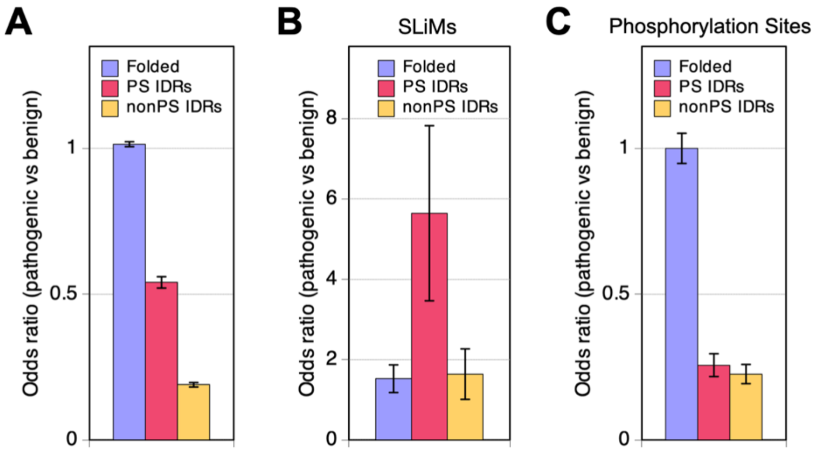

**Figure S3. Pathogenic versus benign odds ratios for missense mutations.** (**A**) By region class, and for mutations found in (**B**) SLiMs or (**C**) at phosphorylation sites, also by region class. Error bars show the standard error.

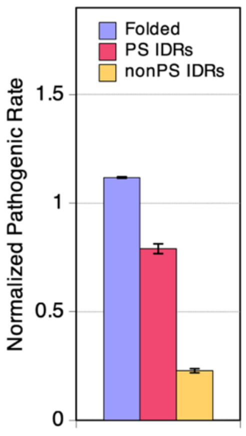

**Figure S4. Pathogenic missense mutation rate by region class for the natural variants index.** Error bars show the standard error.

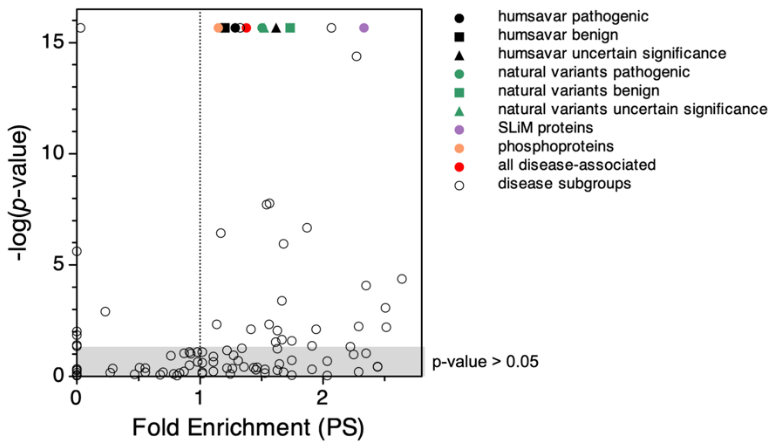

**Figure S5. Enrichment for phase separation in protein sets.** Fold enrichment of phase-separating (PS) proteins and one-tail $p$-values from the Mann-Whitney $U$-test (5) were both calculated using the reference human proteome as the comparison set. The black circle, square and triangle show proteins found in the humsavar index with pathogenic, benign, or uncertain clinical significance missense mutations, respectively. The green circle, square, and triangle show proteins found in the natural variants index with pathogenic, benign, or uncertain clinical significance missense mutations, respectively. Purple and salmon circles show proteins with confirmed SLiMs or confirmed phosphorylation sites, respectively. The red circle shows all proteins annotated as disease-associated in UniProt. Open circles show the disease subgroups from Table 3. Protein sets in the grey shaded region did not exhibit statistically significant differences in predicted enrichment for phase separation when compared to the reference human proteome.

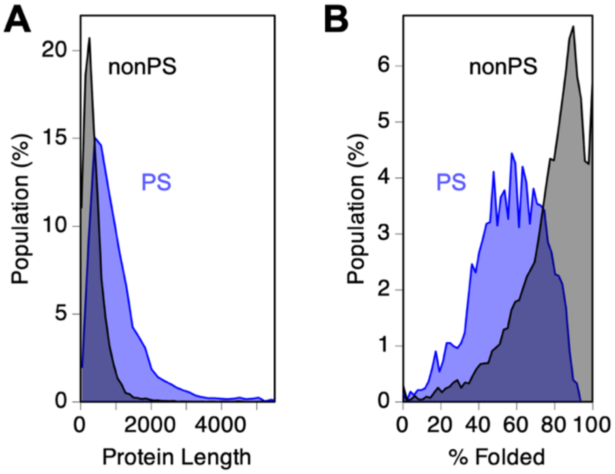

**Figure S6. Protein length and percent of residues folded in PS and nonPS proteins.** Populations were calculated using the reference human proteome. PS proteins had sequence-calculated PS potential ≥100; nonPS proteins had sequence-calculated PS potential <100.

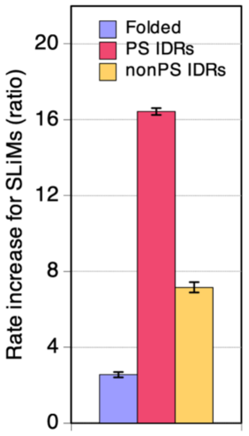

**Figure S7. Pathogenic missense mutation rate increases in SLiMs by region class.** Shown is the ratio of the pathogenic missense mutation rate for SLiM positions (taken from Figure 2A) divided by the pathogenic rate for both SLiM and nonSLiM positions (taken from Figure 1A) by region class. Values >1 indicate the pathogenic mutation rate was higher for SLiM positions in that region. Error bars show the propagated standard error.

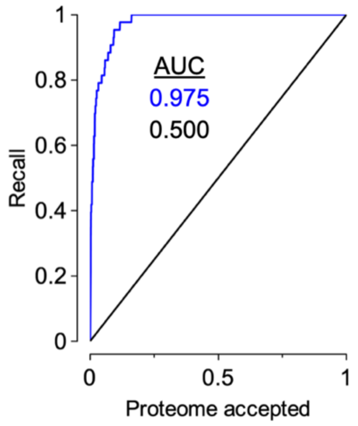

**Figure S8. Quantifying differences in predicted PS potential between protein sets.** For a protein set, the percent of proteins with sequence-calculated PS potential equal to or greater than a given value can be computed, as shown in Figure S2 for the reference human proteome and for confirmed homotypic phase-separating (PS) proteins. Percent of set data can be plotted against the human proteome percent of set, as shown here. The human proteome percent of set plotted against itself gives the identity line (black) and yields 0.5 for its area under the curve (AUC). The homotypic PS protein set (blue) plotted against the human reference proteome yields 0.975 for its AUC.

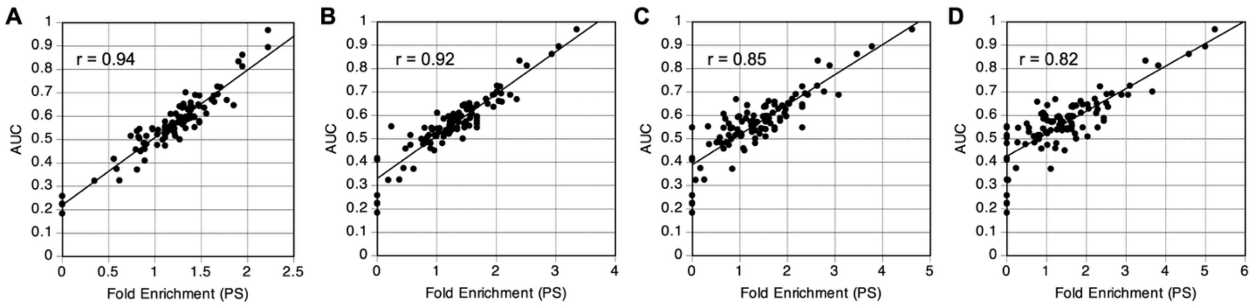

**Figure S9. AUC trends with enrichment for phase separation behavior.** Fold enrichment for PS predicted proteins, relative the reference proteome, was calculated using PS potential cutoffs of (**A**) 25, (**B**) 50, (**C**) 75, and (**D**) 100 for the protein sets listed in Table S3. AUC was calculated using the reference human proteome as the comparison set.

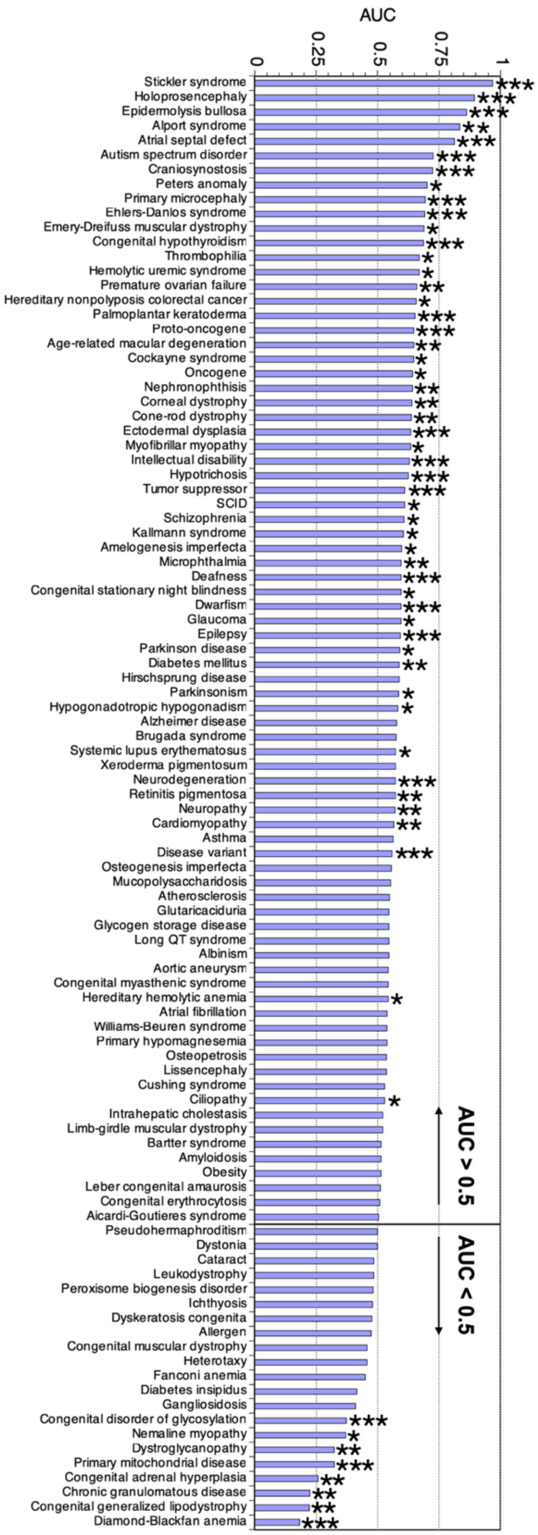

**Figure S10. AUC rank order for protein sets annotated as disease associated.** Curated protein sets confirmed to be associated with human disease were obtained from UniProt (1) and analyzed by ParSe (2). AUC >0.5 predicts enrichment for phase separation behavior. An asterisk marks AUC greater than one standard deviation ($\sigma$) from the mean AUC of similarly sized random protein sets; two asterisks indicate greater than $2\sigma$, and three indicate greater than $3\sigma$.

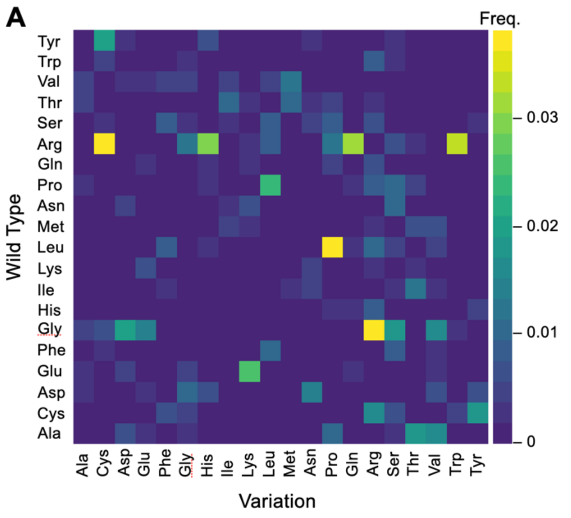

**Figure S11. Frequency of single amino acid missense variation types.** Substitution frequencies (Freq.), representing wild type to variation, were calculated as the number of a type (e.g. Tyr-to-Ala) divided by the total number of single amino acid missense mutations in the humsavar index with (**A**) pathogenic or likely pathogenic and (**B**) benign or likely benign clinical significance. The calculated frequency was colored according to the scale on the right of each plot.

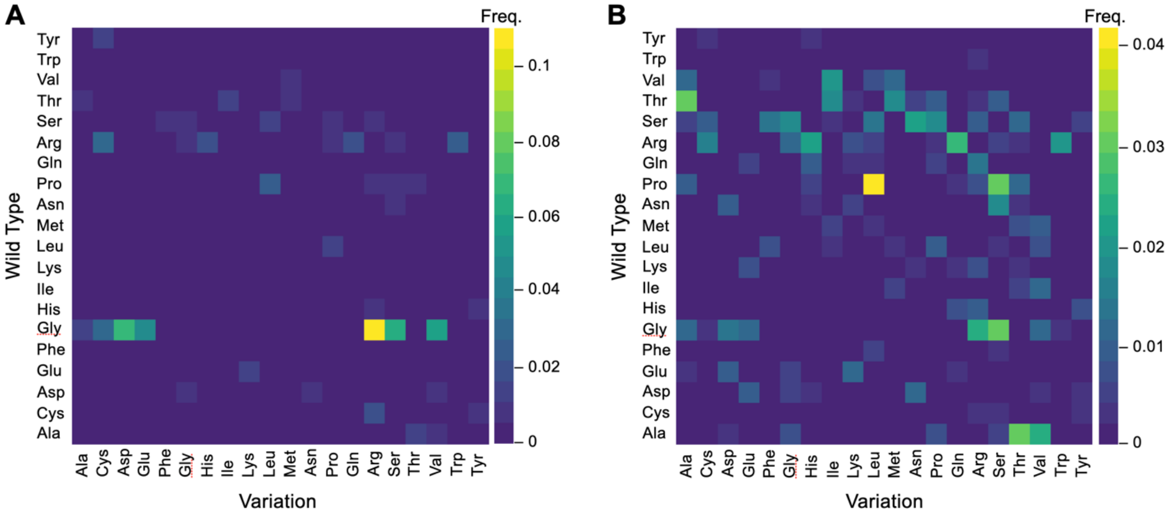

**Figure S12. Frequency of single amino acid missense variation types in PS IDRs.** Substitution frequencies (Freq.) were calculated as the number of a type (e.g. Tyr-to-Ala) found in a PS IDR divided by the total number of single amino acid missense mutations in the humsavar index found in a PS IDR with (**A**) pathogenic or likely pathogenic and (**B**) benign or likely benign clinical significance. The calculated frequency was colored according to the scale on the right of each plot.

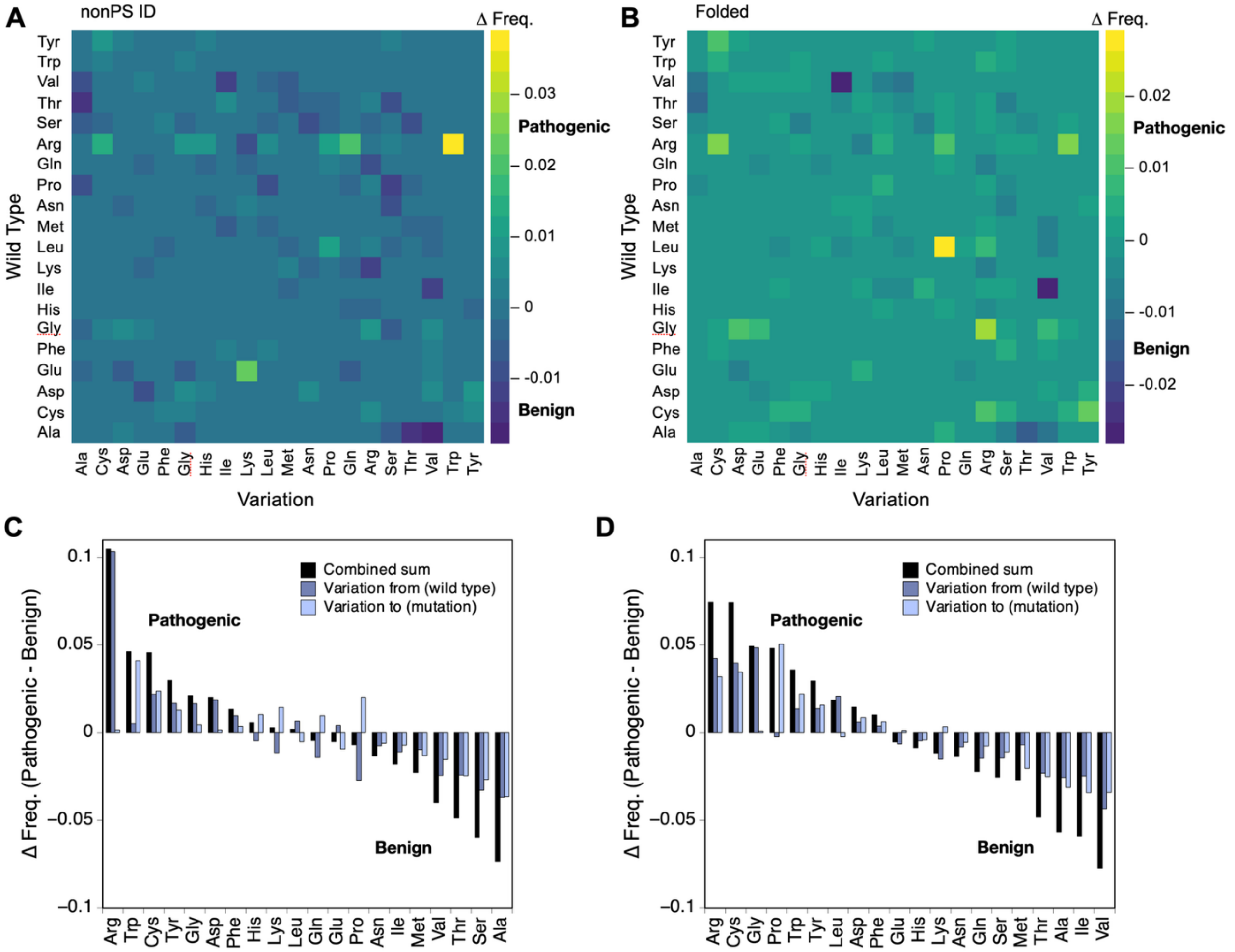

**Figure S13. Pathogenic minus benign difference frequencies for single amino acid missense variation types in nonPS and folded regions.** Difference frequencies (Δ Freq.), colored according to the given scale, were computed from the single amino acid missense variations in the humsavar index for mutations found in regions matching the (**A**) nonPS ID and (**B**) folded classes. Values from (**C**) panel A and (**D**) panel B were summed by wild type amino acid (blue) and mutation amino acid (light blue). The combined sum (black) is the simple addition of the wild type and mutation values.

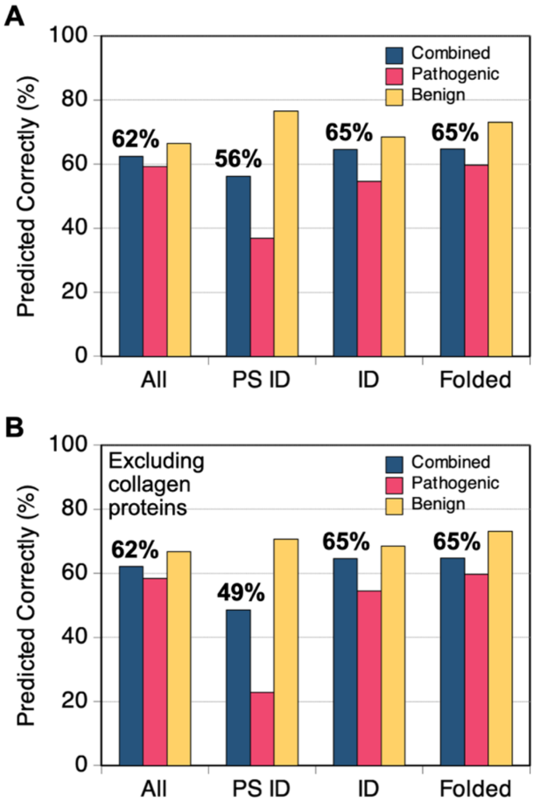

**Figure S14. Predicting clinical significance of missense mutations in the natural variants index.** (**A**) Pathogenic minus benign difference frequencies were calculated for all single amino acid missense mutations in the humsavar index and for the subsets residing in protein regions matching the PS ID, ID (i.e., nonPS ID), or folded classes. Clinical significance was predicted for missense mutations in the natural variants index with known clinical significance where a positive difference frequency predicts pathogenic, and a negative difference frequency predicts benign. Shown is the percent predicted correctly. (**B**) The effect from excluding collagen proteins.
